# Wnt/PCP signaling mediates breast cancer metastasis by promoting pro-invasive protrusion formation in collectively motile leader cells

**DOI:** 10.1101/2022.01.07.475316

**Authors:** Kacey VanderVorst, Courtney A. Dreyer, Jason Hatakeyama, George R. R. Bell, Anastasia L. Berg, Maria Hernandez, Hyun Lee, Sean R. Collins, Kermit L. Carraway

## Abstract

As evidence supporting essential roles for collective cell migration in carcinoma metastasis continues to accumulate, a better understanding of the underlying cellular and molecular mechanisms will be critical to translating these findings to the treatment of advanced cancers. Here we report that Wnt/PCP, a non-canonical Wnt signaling pathway, mediates breast cancer collective migration and metastasis. We observe that mammary gland-specific knockout of *Vangl2*, a tetraspanin-like scaffolding protein required for Wnt5a-induced signaling and motility in cultured breast cancer cell lines, results in a striking decrease in metastatic efficiency but not primary tumor growth in the *MMTV-NDL* transgenic mouse model of HER2-positive breast cancer. We also observe that expression levels of core Wnt/PCP components *Wnt5a*, *Vangl1* and *Vangl2* are selectively elevated in K14-positive leader cells relative to follower cells within a collectively migrating cohort, and that *Vangl2* expression selectively promotes RhoA activation in leading edge cells. Moreover, *Vangl* expression drives collective migration in three-dimensional *ex vivo* tumor organoids, and Vangl protein specifically accumulates within pro-migratory filamentous actin-rich protrusions of leader cells. Together, our observations point to a model whereby Wnt/PCP upregulation facilitates breast tumor collective cell motility by selectively augmenting the formation pro-migratory protrusions within leader cells.

## Introduction

Metastasis is a complex, multi-step process whereby cancer cells invade into surrounding tissues, access and traverse the vasculature, disseminate throughout the body, and proliferate at secondary sites (Talmadge & Fidler, 2010). Observations that carcinoma cells invade almost exclusively in a collective manner (Bronsert et al., 2014), and that metastatic lesions may be largely seeded by polyclonal cell clusters rather than individual disseminated cells (Aceto et al., 2014; Cheung et al., 2016; Fischer et al., 2015; Hou et al., 2012), strongly suggest that collective cell migration, defined as the coordinated movement of cohorts of cells in sheets or clusters that retain cell-cell contacts (Friedl & Gilmour, 2009), is a major driver of invasiveness and metastasis. In non-transformed tissues, collective cell migration promotes blood vessel formation (Geudens & Gerhardt, 2011), convergent extension (Davey & Moens, 2017), branching morphogenesis (Ewald et al., 2008), and wound healing (Friedl & Gilmour, 2009). However, the study of collective cell migration in carcinomas significantly lags that of classical epithelial-mesenchymal transition (EMT)-mediated motility of individual cells. Thus, a better understanding of cell signaling pathways that govern collective cell migration and invasiveness may identify novel therapeutic targets to intervene in patients with aggressive and late-stage disease.

We recently proposed a model whereby aberrant activation of Wnt/planar cell polarity (Wnt/PCP) signaling (VanderVorst et al., 2019), a branch of non-canonical Wnt signaling paradoxically critical to both the establishment and maintenance of polarity in epithelial sheets as well as cell migration during embryonic development (Butler & Wallingford, 2017; Caddy et al., 2010), promotes the invasiveness of primary tumor cells. In Wnt/PCP signaling, binding of non-canonical Wnt ligands such as Wnt5a to transmembrane Frizzled (Fzd) receptors initiates polarization signals that are transduced through the essential pathway components Vangl, Dishevelled (Dvl), and Prickle (Pk) (Chu & Sokol, 2016; Minegishi et al., 2017; Wu et al., 2013). Although Dvl and Fzd are required for both canonical and alternative non-canonical Wnt pathways, the unique engagement of Vangl1 and Vangl2 transmembrane scaffolds in Wnt/PCP signaling may provide the platform necessary for the assembly of pathway-specific complexes (VanderVorst et al., 2018). Vangl1 and Vangl2 are highly similar; their amino acid sequences exhibit 64.3% identity and 78.6% similarity, and no functional biochemical differences have been reported (Hatakeyama et al., 2014). However, Vangl2 alterations result in more profound developmental defects, suggesting a more prominent role for Vangl2 in embryonic tissue organization (Belotti et al., 2012; Hatakeyama et al., 2014). Wnt/PCP signaling is a significant driver of collective cell migration in development (Carmona-Fontaine et al., 2008; Ybot-Gonzalez et al., 2007), and studies employing Looptail (Lp) mice, which harbor point mutations in Vangl2 that alter its trafficking and localization, suggest that Vangl subcellular localization is critical in collectively migrating cells (Murdoch et al., 2001).

Consistent with observations from developmental studies, Wnt/PCP components mediate cell motility in cancer cells (Asad et al., 2014; Kurayoshi et al., 2006), and core Wnt/PCP components are dysregulated in multiple tumor types, including breast (Anastas et al., 2012; Daulat et al., 2016; Luga et al., 2012; MacMillan et al., 2014; Pukrop et al., 2006; Puvirajesinghe et al., 2016; Zhang et al., 2016), brain (Wald et al., 2017), ovarian (Asad et al., 2014), prostate (Uysal-Onganer et al., 2010), gastric (Kurayoshi et al., 2006), and colorectal cancers (Nishioka et al., 2013; Ueno et al., 2008). We have reported that *VANGL1* and *VANGL2* are respectively upregulated in 5% and 24% of invasive breast carcinomas compared to healthy breast tissue (Hatakeyama et al., 2014), and others have found that elevated *VANGL1* and *VANGL2* are also associated with increased recurrence and decreased metastasis-free survival of breast cancer patients (Anastas et al., 2012; Puvirajesinghe et al., 2016). Together, these observations point to the possibility that aberrant Wnt/PCP activation contributes to breast cancer progression by promoting collective cell migration resulting in metastasis.

Here we examine the role of Vangl-dependent Wnt/PCP signaling in breast cancer invasiveness and metastasis. We demonstrate that Vangl2 is critical for efficient metastasis but dispensable for primary tumor growth in ErbB2-induced mouse mammary tumors. We further find that Vangl- dependent Wnt/PCP signaling at the leading edge of migrating breast cancer cells results in increased RhoA GTPase activity and formation of pro-migratory protrusions, resulting in collective cell migration *in vitro* and invasion *ex vivo*.

## Results

### Vangl2 deletion suppresses mammary tumor metastasis to the lungs but does not alter primary tumor growth

We assessed the functional importance of Vangl2 to mammary tumorigenesis and tumor cell metastatic dissemination by specifically ablating *Vangl2* in the mammary epithelium of *MMTV- NDL* mice. In this well-characterized genetically engineered mouse model of breast cancer, an activated ErbB2 mutant encoded by the transgenic rat *c-ErbB2/neu* allele under the control of the MMTV promoter drives the formation of metastatic multifocal mammary tumors at approximately 20 weeks of age (Siegel et al., 1999) (Figure 1A). Effective deletion of *Vangl2* in mammary tumors of *Vangl2^fl/fl^ ;MMTV-Cre^+/-^;MMTV-NDL^+/-^* (Vangl2^fl/fl^/NDL) mice relative to *Vangl2^fl/fl^;MMTV-NDL^+/-^* (Vangl2^+/+^/NDL) mice was confirmed by *q*PCR (Figure 1–figure supplement 1A). Although *Vangl1* may compensate for loss of *Vangl2* in some contexts (Hatakeyama et al., 2014), *Vangl1* transcript is not significantly altered in Vangl2^fl/fl^/NDL mammary tumors relative to Vangl2^+/+^/NDL tumors (Figure 1–figure supplement 1B). *Vangl2* ablation did not produce discernable effects on viability, breeding, or lactation, and no differences in mammary gland architecture were noted between genotypes in adult virgin mammary glands (Figure 1–figure supplement 1C).

**Figure 1.**
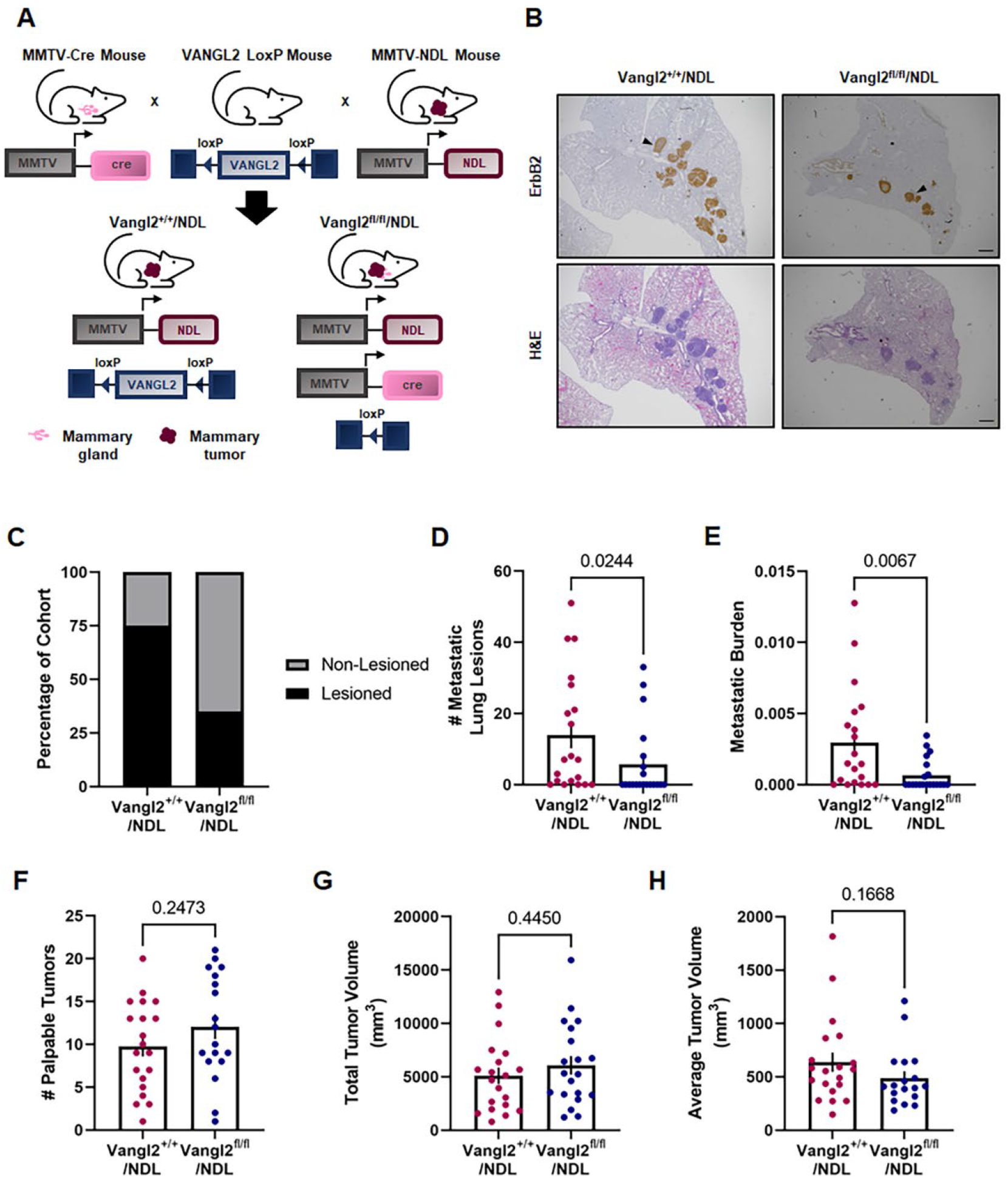
Vangl2 deletion suppresses mammary tumor metastasis to the lung. **A** Summary of the transgenic mouse strategy employed to assess Vangl2 deletion in mammary tumorigenesis. **B** Representative images of formalin fixed, paraffin embedded sections from Vangl2^+/+^/NDL and Vangl2^fl/fl^/NDL lung tissue following immunodetection of ErbB2 (top panel) and H&E staining (bottom panel). Examples of ErbB2-positive metastatic lung lesions are denoted by black arrowheads, scale bar =500µm. **C-E** Lung lobes (5 lobes per mouse) were evaluated by histology for the occurrence of metastatic lesions for Vangl2^+/+^/NDL (*n*=20) and Vangl2^fl/fl^/NDL tumor-bearing mice (*n*=20). The number of mice bearing metastatic lesions **(C)**, numbers of metastatic lesions **(D)**, and metastatic burden **(E)** were assessed. **F-H** Vangl2/NDL primary tumor growth characteristics were assessed for Vangl2^+/+^/NDL (*n*=20) and Vangl2^fl/fl^/NDL (*n*=20) tumor-bearing animals, including number of palpable tumors **(F)**, total tumor volume **(G)**, and average tumor volume **(H)**. Significance was determined by Mann-Whitney test and bar graphs represent the mean ± sem of experimental replicates (*n*).

Despite similar kinetics of primary tumor initiation and growth in Vangl2^+/+^/NDL and Vangl2^fl/fl^/NDL mice (Figure 1–figure supplement 2A-D), *Vangl2*-depleted tumors are significantly less metastatic than *Vangl2*-intact tumors (Figure 1B-E). Analysis of lung tissue revealed that deletion of *Vangl2* in *MMTV-NDL* tumors results in significantly reduced frequency of metastatic disease (Figure 1C), number of lung metastases (Figure 1D) and overall metastatic burden (Figure 1E) despite similar primary tumor characteristics such as numbers of palpable tumors (Figure 1F), total tumor volume (Figure 1G), average tumor volume (Figure 1H), tissue histology (Figure 1–figure supplement 2E,F), proliferative capacity (Figure 1–figure supplement 2G,H), and apoptosis (Figure 1–figure supplement 2I,J). Importantly, *Vangl2* appears to be critical to successful metastatic colonization of the lungs (Figure 1D,E) but is not required for proliferation of metastatic lesions in Vangl2^+/+^/NDL and Vangl2^fl/fl^/NDL mice (Figure 1–figure supplement 3A,B). Further, cells derived from Vangl2^+/+^/NDL and Vangl2^fl/fl^/NDL tumors injected into the tail veins of FvB/NJ mice exhibit no differences in metastatic lesion colonization efficiency (Figure 1–figure supplement 3C-F). Taken together, these findings suggest that the reduced incidence of metastasis observed upon *Vangl2* ablation is the result of reduced dissemination from the primary tumor rather than suppressed outgrowth of colonies in the lungs. Because Wnt/PCP signaling is vital to collective cell motility events critical to embryonic tissue patterning (Butler & Wallingford, 2017; Hatakeyama et al., 2014), we hypothesized that Vangl2 facilitates local invasion and migration from the primary tumor, resulting in dissemination and metastatic disease.

### Vangl2-dependent Wnt/PCP signaling promotes breast cancer cell migration

We interrogated whether loss of Vangl2 impacts breast cancer cell motility by employing a panel of human breast cancer cell lines encompassing several molecular subtypes: triple-negative BT549, HCC1937, HCC38, MDA-MB-157, MDA-MB-231 and MDA-MB-468, and ER/PR-positive MCF7. Cells were transduced with lentivirus encoding *VANGL2*-targeted shRNAs and knockdown was confimed by *q*PCR (Figure 2–figure supplement 1A). Loss of *VANGL2* expression significantly impairs breast cancer cell migration, indicated by the reduced ability of *VANGL2* knockdown cells to migrate into a scratch made in the cellular monolayer (Figure 2A), but does not siginificantly impair cell proliferation after 12 hours (Figure 2–figure supplement 1B). These data demonstrate that Vangl2 is critical to breast cancer cell motility, regardless of breast cancer subtype.

**Figure 2.**
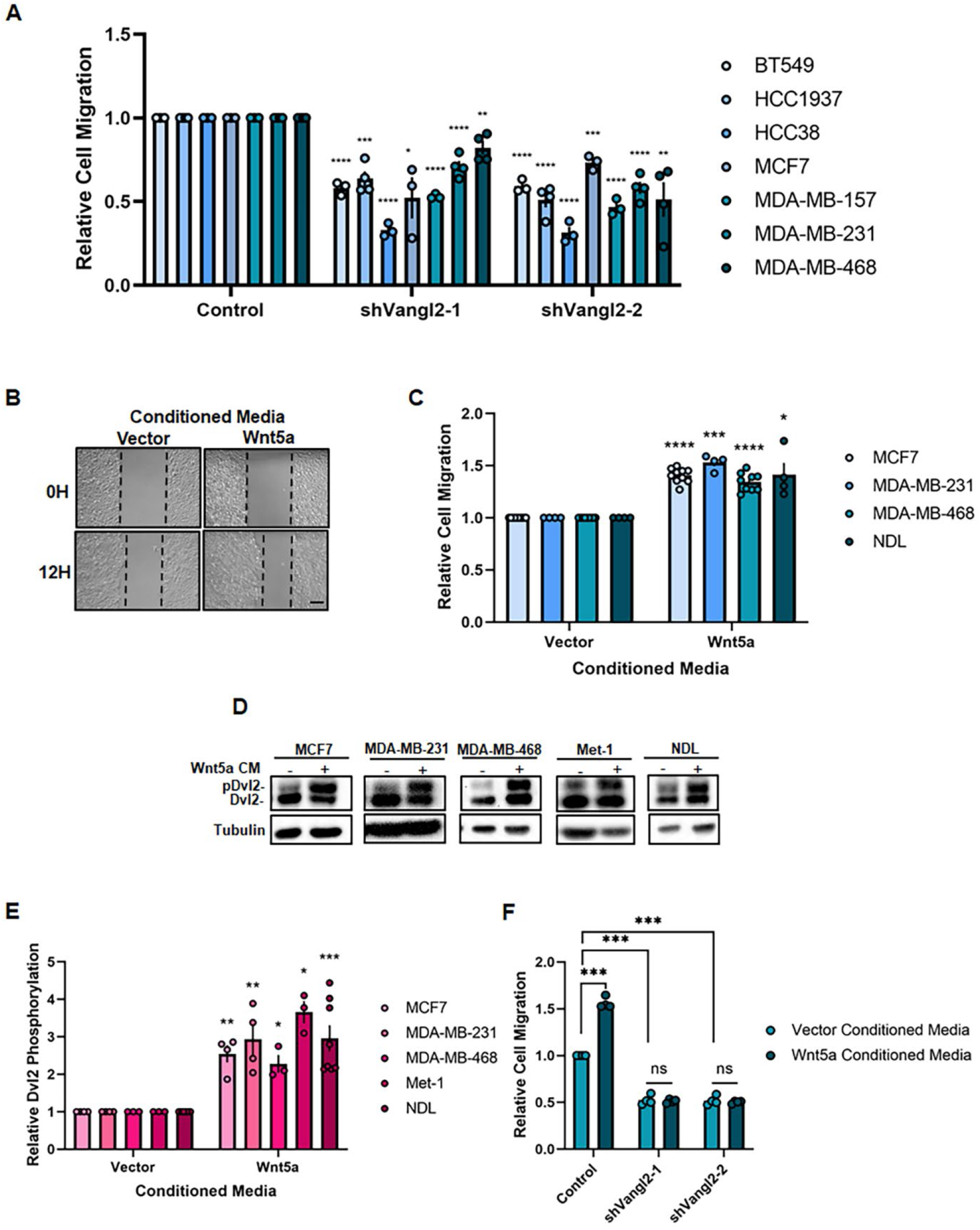
Vangl2 is essential for Wnt5a-induced Wnt/PCP signaling and breast cancer cell motility. **A** Relative cell migration quantification of BT549, HCC1937, HCC38, MCF7, MDA-MB- 157, MDA-MB-231, and MDA-MB-468 cells stably expressing Control, shVangl2-1, or shVangl2- 2 (BT549 *n*=3, *p*=4.81E-05 and *p*=4.47E-05, HCC1937 *n*=4, *p*=0.0002 and *p*=3.37E-05, HCC38 *n*=3, *p*=7.69E-06 and *p*=4.31E-05, MCF7 *n*=3, *p*=0.0180 and *p*=0.0004, MDA-MB-231 *n*=4, *p*=0.0001 and *p*=2.73E-05, MDA-MB-468 *n*=4, *p*=0.0046 and *p*=0.0033). **B** Representative images of migrating NDL cells stimulated with Vector- or Wnt5a- conditioned media at 0 and 12 hours. Scale bar = 200µm. **C** Relative cell migration quantification of MCF7, MDA-MB-231, MDA-MB-468, and NDL cells stimulated with Vector- or Wnt5a-conditioned media (MCF7 *n*=10, *p*=1.54E-08, MDA-MB-231 *n*=4, *p*=0.0006, MDA-MB-468 *n*=9, *p*=2.33E-06, NDL *n*=4, *p*=0.0344,). **D,E** MCF7, MDA-MB-231, MDA-MB-468, Met-1, and NDL cells stimulated with Vector- or Wnt5a- conditioned media for 1 hour blotted for Dvl2 **(D)** and quantification of relative Dvl2 phosphorylation (MCF7 *n*=4, *p*=0.0067, MDA-MB-231 *n*=4, *p*=0.0199, MDA-MB-468 *n*=3, *p*=0.0327, Met-1 *n*=3, *p*=0.0121, NDL *n*=8, *p*=0.0007) **(E)**. **F** Relative cell migration quantification of MDA-MB-231 cells stably expressing Control, shVangl2-1, or shVangl2-2 stimulated with Vector- or Wnt5a- conditioned media (control: vector- vs Wnt5a-conditioned media *n*=4, *p*=0.0003, Control vs shVangl2-1 + vector-conditioned media *n*=4, *p*=0.0003, control vs shVangl2- 2 + vector-conditioned media *n*=4, *p*= 0.0003, shVangl2-1: vector- vs Wnt5a-conditioned media *n*=4, *p*=0.7256, shVangl2-2: vector- vs Wnt5a-conditioned media *n*=4, *p*=0.5804). Bar graphs represent the mean ± sem of experimental replicates (*n*). Significance was determined by a two-sided unpaired *t*-test with Welch’s correction, \**p <* 0.05, \*\**p <* 0.01, \*\*\**p* < 0.001, \*\*\*\**p* < 0.0001.

In motile cells, activation of Vangl-dependent Wnt/PCP signaling occurs by binding of a non-canonical Wnt ligand such as Wnt5a to Fzd receptors at the plasma membrane, resulting in recruitment and phosphorylation of Dvl. Transmembrane proteins Vangl1 and Vangl2 and activated Dvl may serve as both scaffolds and activators of downstream effector components that mediate context- and tissue-specific actin cytoskeletal rearrangements to promote cellular motility (Wald et al., 2017). Consistent with previous reports (MacMillan et al., 2014), we found that Wnt5a is a potent activator of Wnt/PCP signaling that drives breast cancer cell migration. Stimulation of breast cancer cell lines with Wnt5a-conditioned media enhances cellular migration (Figure 2B,C) and robustly increases phosphorylation of Dvl2 (Figure 2D,E) compared to vector control-conditioned media. To exclude the possibility that Wnt5a activates the canonical Wnt pathway, we assessed phosphorylation of β-catenin, a marker of canonical Wnt signaling (van Amerongen, 2012). Stimulation of MCF7 cells with Wnt5a-conditioned media does not significantly impact β- catenin phosphorylation, whereas stimulation with conditioned media containing the potent canonical Wnt activating ligand Wnt3a significantly reduces β-catenin phosphorylation (Figure 2–figure supplement 1C-D). Thus, Wnt5a-dependent migration and Dvl2 phosphorylation in breast cancer cells is driven by engagement of a non-canonical rather than canonical Wnt signaling pathway.

Dvl2 phosphorylation is also a common feature of other non-canonical Wnt pathways independent of Wnt/PCP signaling (Semenov et al., 2007), and while a downstream effector specific to Wnt/PCP signaling has yet to be identified, this branch of non-canonical Wnt signaling requires the formation of Vangl-dependent complexes (Hatakeyama et al., 2014). We determined that Wnt5a-mediated migration is ablated in *VANGL2* knockdown breast cancer cells (Figure 2F), demonstrating that Wnt5a specifically activates Vangl-dependent Wnt/PCP signaling in breast cancer cells.

### High Vangl expression aberrantly engages Wnt/PCP signaling and enhances breast cancer cell motilty

Our observations that Vangl2 mediates mammary tumor metastasis (Figure 1) and is required for Wnt5a-mediated cell migration (Figure 2), combined with previous reports that elevated *VANGL2* expression correlates with worsened metastasis-free survival in breast cancer patients (Puvirajesinghe et al., 2016), suggest that high Vangl expression may result in enhanced cellular migration and aberrant engagement of Wnt/PCP signaling in breast tumors. To investigate this possibility, we stably overexpressed Vangl1 or Vangl2 via lentiviral infection in human and mouse breast tumor cell lines (Figure 3–figure supplement 1A,B). Breast cancer cells overexpressing Vangl1 or Vangl2 exhibit increased motility and Dvl2 phosphorylation compared to cells transduced with control lentivirus (Figure 3A, C-F), and display a distinctive hyper-protrusive leading edge morphology (Figure 3B). These observations indicate that Vangl proteins are sufficient to aberrantly engage Wnt/PCP signaling either independent of Wnt ligand or by potentiating signaling from endogenous Wnt ligand.

**Figure 3.**
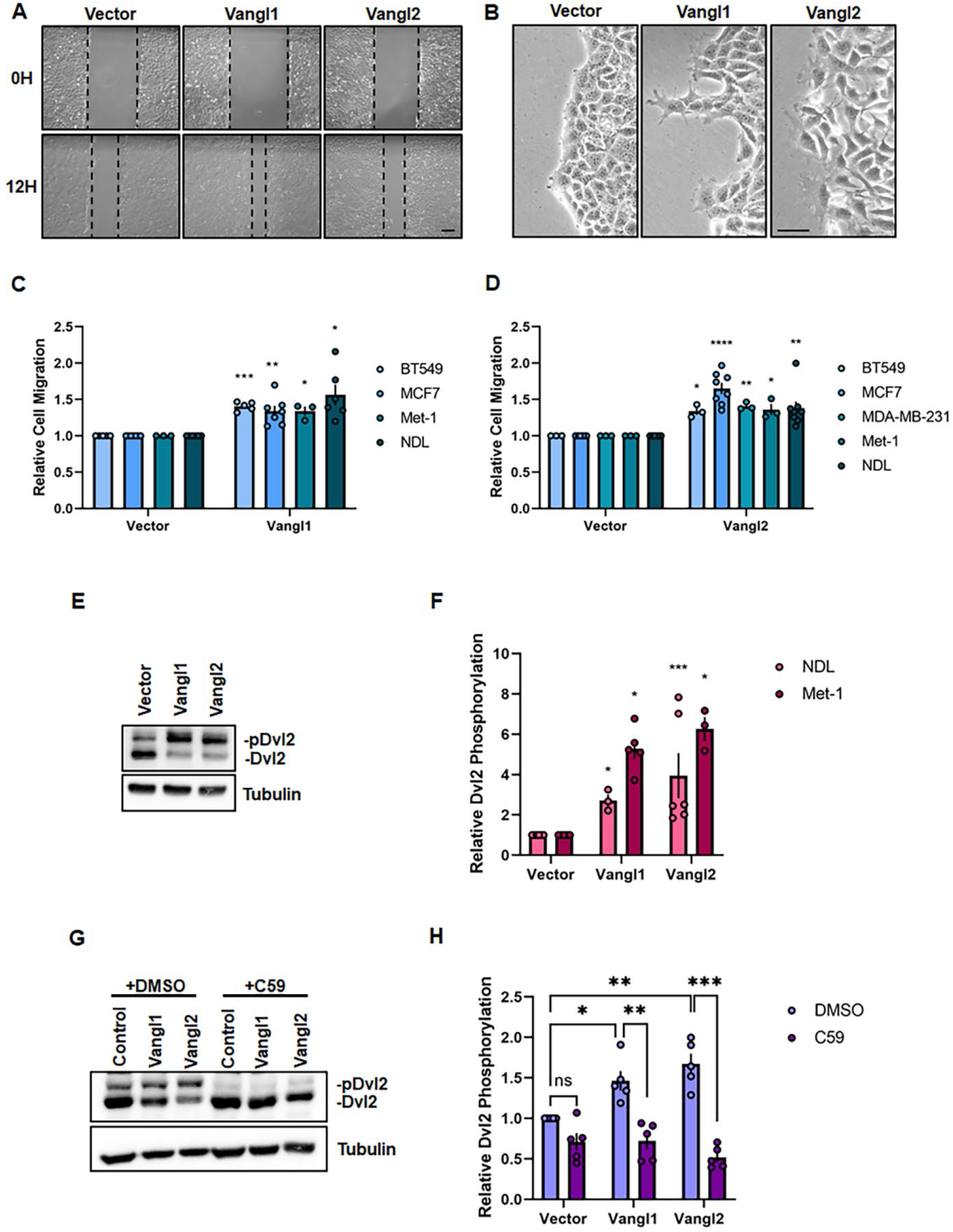
Vangl overexpression potentiates Wnt/PCP signaling and promotes breast cancer cell migration. **A,B** Representative brightfield images of migrating NDL cells stably expressing Vector, Vangl1, or Vangl2 at 0 and 12 hours, scale bar = 200µm **(A)** and leading-edge dynamics at 12 hours, scale bar = 50 µm **(B)**. **C** Relative cell migration quantification of BT549, MCF7, Met-1, and NDL cells stably expressing Vector or Vangl1 (BT549 *n*=5, *p*=0.0001, MCF7 *n*=7, *p*=0.0035, Met-1 *n*=3, *p*=0.0339, NDL *n*=6, *p*=0.0108). **D** Relative cell migration quantification of BT549, MCF7, MDA-MB-231, Met-1, and NDL cells stably expressing Vector or Vangl2 (BT549 *n*=3, *p*=0.0210, MCF7 *n*=8, *p*=6.17E-05, MDA-MB-231 *n*=3, *p*=0.0050, Met-1 *n*=3, *p*=0.0412, NDL *n*=8, *p*=0.0049). **E,F** NDL cells stably expressing Vector, Vangl1 or Vangl2 blotted for Dvl2 **(E)** and quantification of Dvl2 phosphorylation in NDL and Met-1 cells stably expressing Vector, Vangl1 or Vangl2 **(F)** (NDL-Vangl1 *n*=3, *p*=0.0291, NDL-Vangl2 *n*=6, *p*=0.0453, Met-1- Vangl1 *n*=5, *p*=0.0009, Met-1-Vangl1 *n*=3, *p*=0.0119). **G,H** NDL cells stably expressing Vector, Vangl1, or Vangl2 treated with DMSO or 100nM C59 for 24 hours blotted for Dvl2 **(G)** and quantification of relative Dvl2 phosphorylation **(H)**. Bar graphs represent the mean ± sem of experimental replicates (*n*). Significance was determined by a two-sided unpaired *t*-test with Welch’s correction, \**p <* 0.05, \*\**p <* 0.01, \*\*\**p* < 0.001, \*\*\*\**p* < 0.0001.

To distinguish between these possibilities, we treated Vangl overexpressing breast cancer cells with the Porcupine antagonist C59, which impairs palmitoylation and subsequent secretion of Wnt ligands (Proffitt et al., 2013), to deplete endogenous Wnt5a ligand. C59 treatment resulted in ablation of Vangl-mediated Dvl2 phosphorylation (Figure 3G,H), demonstrating that aberrant Wnt/PCP signaling mediated by Vangl overexpression is Wnt ligand-dependent. Taken together, these findings suggest that activation of Wnt/PCP signaling in breast cancer cells, accomplished either by exposing cells to elevated levels of non-canonical Wnt ligand or by potentiating signals from endogenous Wnt ligands through overexpression of Vangls, enhances cellular motility and may promote metastatic dissemination of primary tumor cells.

### Wnt/PCP signaling drives breast carcinoma cell collective invasion

Collective cell invasion is driven by leader cells that aggressively invade while remaining attached to follower cells, resulting in the formation of contiguous invasive strands (Cheung et al., 2013; Cheung et al., 2016; VanderVorst et al., 2019). Invasive leader cells are molecularly and behaviorally distinct from bulk tumor cells, and in some mammary tumor models and human breast tumors express the basal epithelial marker cytokeratin 14 (K14) (Cheung et al., 2013). Importantly, K14-positive leader cells are not enriched for markers of stemness or EMT (Cheung et al., 2013), underscoring the unique character of this population and distinguishing these cells from classically defined invasive mediators of metastasis.

To investigate the contribution of Wnt/PCP signaling to breast cancer cell collective invasion, we employed an *ex vivo* 3D collagen invasion assay (Nguyen-Ngoc et al., 2012), in which tumor organoids form multicellular epithelial cell protrusions that invade a collagen matrix. Individual tumor cells derived from the highly aggressive, metastatic *MMTV-PyMT* mouse model (Guy et al., 1992; Lin et al., 2003) were first seeded in Matrigel, transduced with lentivirus encoding Wnt/PCP components (Figure 4–figure supplement 1A-C), and cultured for one week to generate tumor organoids. Organoids were then transferred to 3D collagen I gels, a model for the microenvironment surrounding invasive breast cancers (Nguyen-Ngoc et al., 2012), and a fraction of epithelial cells became K14-positive leader cells that formed multicellular protrusions of collectively invading cells upon stimulation with *b*FGF (Cheung et al., 2013).

We observed that lentiviral-mediated overexpression of Wnt5a, Vangl1, or Vangl2 significantly increases the frequency of *b*FGF-dependent collectively invading strands formed by *MMTV-PyMT* organoids (Figure 4A-C). Expression of Wnt/PCP components was not sufficient to stimulate collective invasion in the absence of *b*FGF (Figure 4–figure supplement 1D), indicating that Wnt/PCP signaling cooperates with additional signaling pathways to promote collective invasion rather than independently driving formation of invasive protrusions. Our data suggest that Vangl1 is highly expressed in K14-positive cells at the tip of invading strands compared to K14-negative follower cells or non-invading tumor organoid cells (Figure 4D), indicating that Wnt/PCP signaling specifically augments the protrusive activity of K14-positive leader cells that drive collective invasion. In support of our findings, analysis of a publicly available dataset that accompanied the foundational study describing the contributions of K14-positive leader cells to breast cancer progression (Cheung et al., 2016) revealed that *Wnt5a*, *Vangl1,* and *Vangl2* transcripts are significantly elevated in the K14-positive tumor cell population (Figure 4E). Other Wnt/PCP component transcripts including noncanonical Frizzled receptors and Dvl were not significantly altered (Figure 4–figure supplement 1E). Taken together, these findings suggest that Wnt/PCP signaling may augment the invasive behavior of K14-positive leader cells and supports a model in which Vangl-mediated Wnt/PCP signaling drives metastatic dissemination through promotion of local collective cell migration and invasion from the primary tumor.

**Figure 4.**
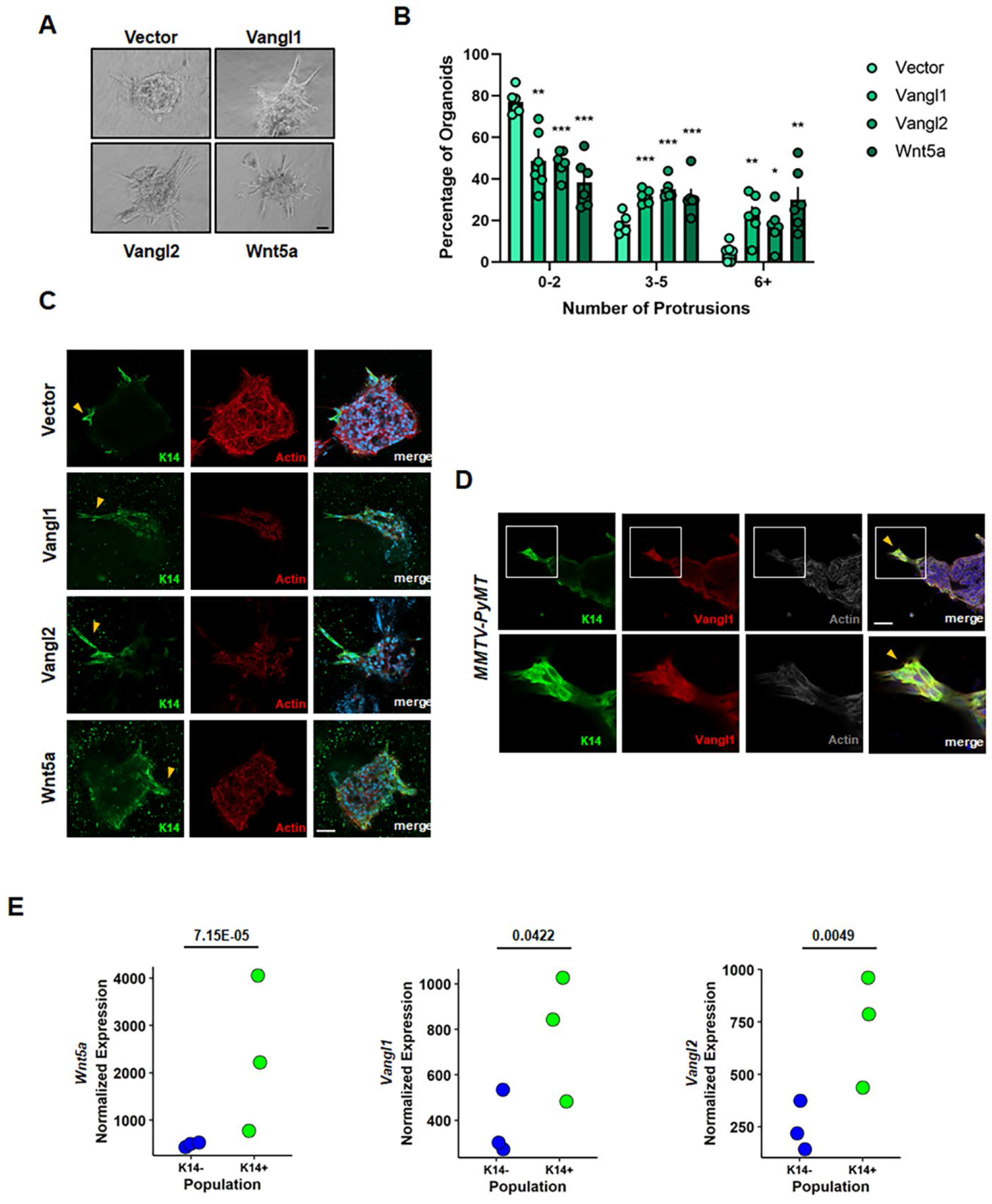
Wnt/PCP signaling drives collective cell invasion *ex vivo* and is upregulated in the K14-positive leader cell population. **A** Representative bright field images of *MMTV-PyMT*- derived mouse mammary tumor organoids stably overexpressing Vector, Vangl1, Vangl2 or Wnt5a invading into collagen in the presence of 2.5nm *b*FGF. **B** Quantification of the percentage of organoids counted with 0-2, 3-5, and 6+ collectively invading protrusions for Vector-, Vangl1-, Vangl2-, or Wnt5a-expressing organoids (Vector 547, Vangl1 371, Vangl2 276, Wnt5a 456 organoids counted from *n*=6 independent experiments, *p*-values represent Vector (V) vs Vangl1 (V1), Vangl2 (V2), Wnt5a (W), 0-2 protrusions; V vs V1 *p*=0.0011, V vs V2 *p*=6.32E-06, V vs W *p*=2.50E-05, 3-5 protrusions; V vs V1 *p*=0.0001, V vs V2 *p*=8.94E-05, V vs W *p*=0.0092, 6+protrusions; V vs V1 *p*=0.0027, V vs V2 *p*=0.0132, V vs W *p*=0.0024). Bar graphs represent the mean ± sem of experimental replicates (*n*), significance was determined by a two-sided unpaired *t*-test, \**p <* 0.05, \*\**p <* 0.01, \*\*\**p* < 0.001, \*\*\*\**p* < 0.0001. **C** Representative confocal images of Vector-, Vangl1-, Vangl2-, or Wnt5a-expressing PyMT-derived organoids stained with K14: green, Actin: red, and DAPI: blue. Yellow arrows: K14+ collectively invading protrusion. **D** Representative confocal images of PyMT-derived organoids stained with K14: green, Vangl1: red, Actin: grey, and DAPI: blue (top) and boxed regions have been expanded to show details (bottom). Yellow arrows: K14+/Vangl1+ leader cells of a collectively invading protrusion. **E** Analysis of RNA-sequencing data set SRP066316 from NCBI Sequence Read Archive for *Wnt5a, Vangl2,* and *Vangl1* transcript in K14- and K14+ cells derived from *MMTV-PyMT* tumors (*Wnt5a p*= 7.15E-05, *Vangl2 p*=0.0049, *VANGL1 p*=0.0422), significance was determined by likelihood ratio test followed by Benjamin-Hochberg correction for multiple hypothesis testing. Scale bars=50µm.

### Vangl localizes to the leading-edge of collectively migrating breast cancer cells and promotes a hyper-protrusive leading-edge

Wnt/PCP signaling is essential for the establishment and maintenance of polarity in epithelial tissues, where it regulates cell polarization in the planar axis across the surface of an epithelial sheet (Devenport, 2014). Planar polarity across the tissue is achieved through the asymmetric distribution of core Wnt/PCP complexes within individual cells reinforced by intracellular antagonism between opposing complexes (Axelrod, 2001; Jenny et al., 2005; Tree et al., 2002; Warrington et al., 2017). Intercellular complexes formed by opposing complexes on adjacent cells then transmit that asymmetry to neighboring cells (Chen et al., 2008; Strutt & Strutt, 2008; Wu & Mlodzik, 2008), and propagation of this asymmetry across many cell distances allows for integration of global cues to locally polarized cellular behavior (Devenport, 2014). However, the requirement for Wnt/PCP complex asymmetry in migrating cancer cells has remained unclear, despite significant effort to understand component localization in migrating breast cancer cells (Anastas et al., 2012; Daulat et al., 2016; Luga et al., 2012).

We employed immunofluorescence microscopy to assess the localization of endogenous Wnt/PCP components in MCF7 breast cancer cells and cells derived from *MMTV-PyMT* tumors, which migrate as cohesive sheets with E-cadherin-rich cell-cell junctions (Figure 5–figure supplement 1A,B). In MCF7 cells, we observed that both Vangl1 and Fzd7, which typically localize to opposing complexes within planar polarized tissues (VanderVorst et al., 2018), co-localize at actin-rich migratory protrusions in cells at the leading-edge of a collectively migrating cohort (Figure 5A). Consistent with observations that Vangl overexpression enhances cellular motility (Figure 3C,D), Vangl1 overexpression in MCF7 cells elicits a hyper-protrusive leading-edge enriched for Vangl1, Fzd7 and Actin (Figure 5B), suggesting that elevated Vangl1 mediates the assembly of Wnt/PCP complexes that promote the formation of pro-migratory protrusions that drive collective cell migration. Indeed, elevated Vangl1 expression significantly increases both the number of Vangl1-rich protrusions in leader cells and the percentage of leader cells with Vangl1- rich protrusions in a collectively migrating sheet of MCF7 cells (Figure 5–figure supplement 1C,D). We also observed Vangl1-rich protrusions at the leading-edge of collectively migrating primary *MMTV-PyMT* tumor cells migrating as both sheets and clusters (Figure 5C). These data suggest that Vangl mediates the formation of pro-migratory protrusions in collectively migrating breast cancer cells and that high Vangl expression drives enhanced cellular invasiveness through the regulation of aberrant leading-edge protrusion formation.

**Figure 5.**
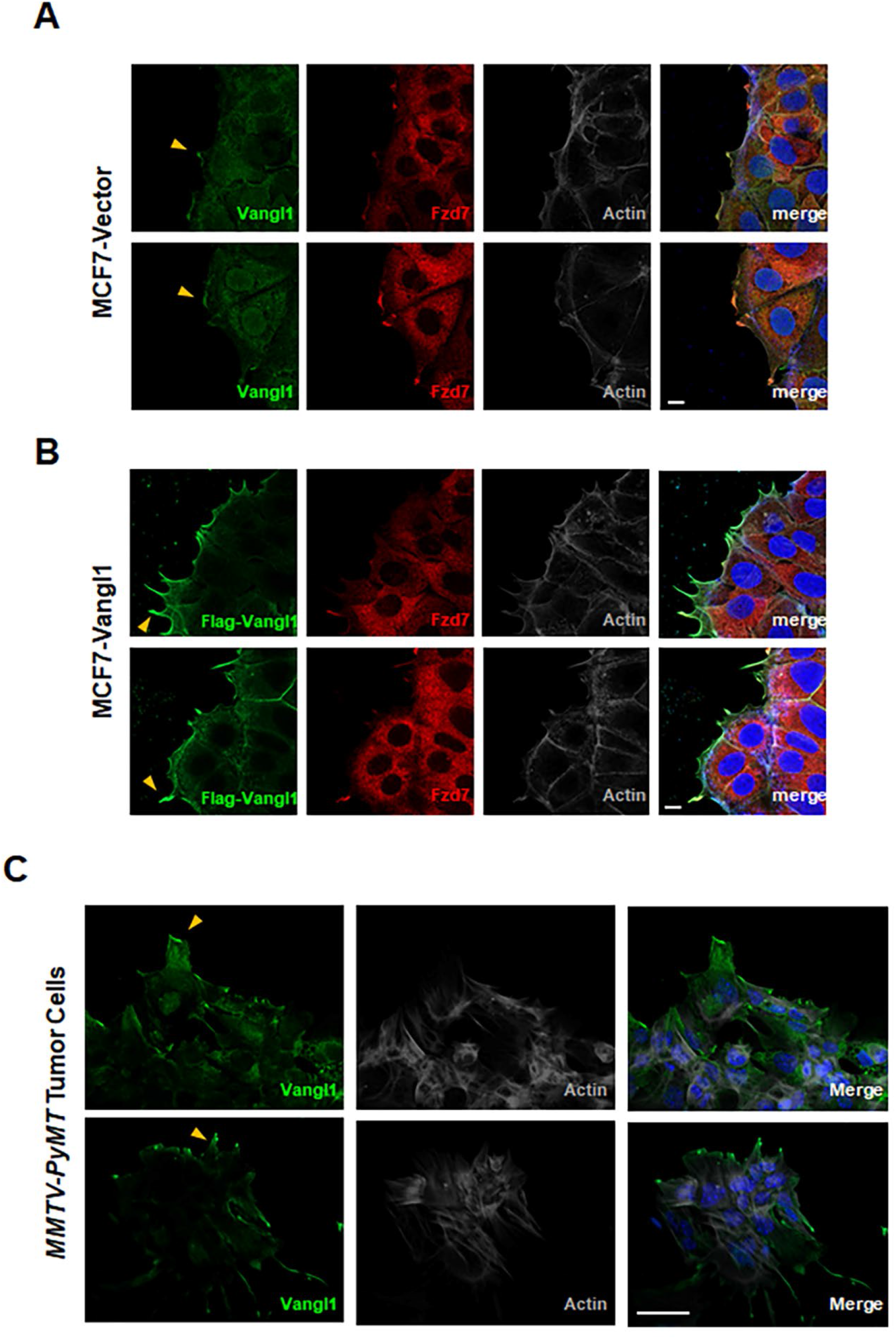
Vangl localizes to the leading-edge of leader cells in collectively migrating breast cancer cells and promotes a hyper-protrusive morphology. A,B Representative confocal images of collectively migrating MCF7-Vector cells stained for Vangl1: green, Fzd7: red, Actin: grey, and DAPI: blue (A) and MCF7-Flag-Vangl1 cells stained for Flag: green, Fzd7: red, Actin: grey, and DAPI: blue (B), yellow arrows: Vangl1-rich protrusions, scale bar=10µm. C Representative confocal images of *MMTV-PyMT* tumor-derived cells migrating collectively as a sheet (upper panel) or as a cluster (lower panel) stained for Vangl1: green, Actin: grey, and DAPI: blue. Yellow arrows: Vangl1-rich protrusions in leading-edge cells of migrating sheets and clusters, scale bar=50µm.

### Vangl2 regulates RhoA activity in leader cells of collectively migrating breast cancer cells

Our findings that Vangl drives collective cell motility and invasion as well as mediates the formation of pro-migratory protrusions in leader cells of collectively migrating breast cancer cells led us to question the molecular mechanisms by which Vangl achieves these outcomes. We hypothesized that Vangl may regulate the actin cytoskeleton in leader cells via engagement of Rho GTPases Rac1 and RhoA, which are engaged in Wnt/PCP-mediated motility during vertebrate gastrulation in developing embryos (Habas et al., 2003; Habas et al., 2001). The regulation of Rac1 and RhoA GTPase activity is complex and permits context-specific activation of signaling events at specific subcellular localizations with precise kinetics (Ridley, 2015). Unfortunately, previous studies that investigated the ability of Wnt/PCP components to specifically engage and regulate Rho GTPases in cancer cells have predominantly assessed global GTPase activity in lysed cells via GST pull-down assays (Asad et al., 2014; Kurayoshi et al., 2006; Wald et al., 2017), leaving the localization and kinetics of Wnt/PCP-mediated RhoA and Rac1 activity largely unexplored.

We investigated the spatiotemporal dynamics of GTPase signaling in real time via time-lapse imaging in collectively migrating breast cancer cells by monitoring GTPase activity using stably expressed Rac1 or RhoA fluorescence resonance energy transfer (FRET) biosensors (Yang et al., 2016). Here, MCF7 cells stably expressing non-targeting control shRNA or two independent *VANGL2*-targeted shRNAs and a Rac1 or RhoA biosensor were seeded onto glass-bottom plates, the confluent monolayer was scratched at zero hours, and scratches imaged every fifteen minutes throughout the 12 hour migration assay (Figure 6A, Supplementary Videos 1-3). The mean FRET ratio, which indicates Rac1 or RhoA activity, was measured after 1, 6, and 12 hours of migration and plotted as a function of distance from the leading-edge of the migrating cohort of MCF7 cells using a custom MATLAB script to quantify Rac1 and RhoA activity across the monolayer of cells (Figure 6B, Figure 6–figure supplement 1A). Briefly, our MATLAB script identified the leading-edge of the migrating cohort of MCF7 cells (Figure 6–figure supplement 1B) and binned migrating cells based on their distance from the edge of the scratch (Figure 6–figure supplement 1C).

**Figure 6.**
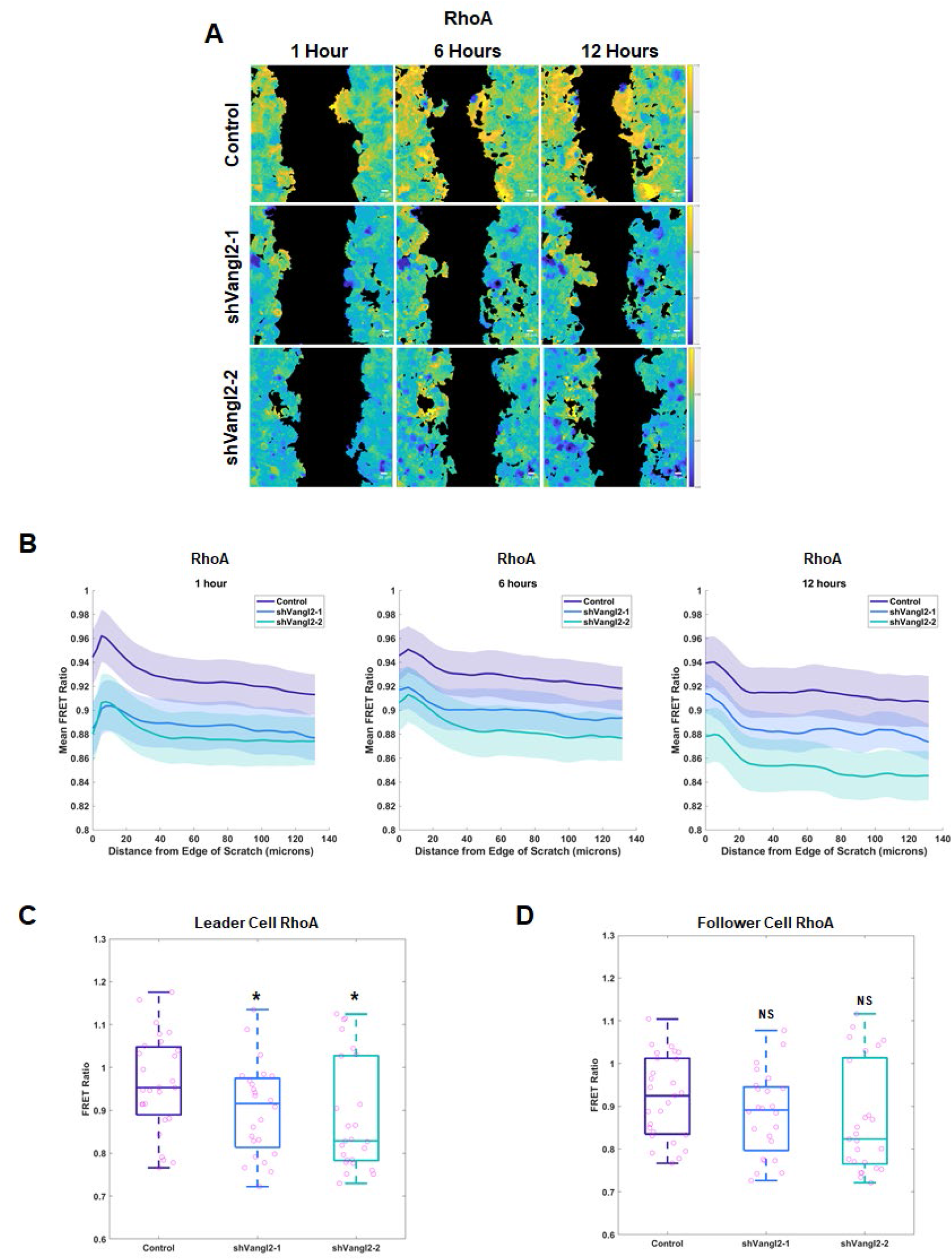
Vangl2 preferentially regulates RhoA activity in leader cells of collectively migrating breast cancer cells. **A** Representative spatial activity profiles of RhoA in collectively migrating MCF7 cells stably expressing RhoA-FRET biosensor and Control (top row), shVangl2- 1 (middle row), or shVangl2-2 (bottom row) at 1 hour (left column), 6 hours (center column), and 12 hours (right column. Color bars indicate the range of RhoA-FRET biosensor ratios. Scale bar=25µm. **B** RhoA activity as a function of distance in µm from the leading-edge of collectively migrating MCF7 cells stably expressing Vector (*n*=27 wells), shVangl2-1 (*n*=24 wells), or shVangl2-2 (*n*=25 wells) at 1, 6, and 12 hours of migration, error bars indicate ± sem. **C,D** RhoA activity after one hour of migration at 5µm **(C)** and 100µm **(D)** from the leading-edge of collectively migrating MCF7 cells stably expressing Vector, shVangl2-1 (Vector vs shVangl2-1, 5µm *p*=0.05, 100µm *p*=0.1803), or shVangl2-2 (Vector vs shVangl2-2, 5µm *p*=0.0271, 100µm *p*=0.1094), significance was determined by a two-sided unpaired *t*-test.

Spatial analysis of RhoA activity after one hour of migration revealed that RhoA activity is highest in the leading-edge cells, with activity peaking at approximately 5-10µm from edge of the scratch (Figure 6B) in a collectively migrating cohort of MCF7 cells. Indeed, RhoA activity is significantly higher 5µm from the edge of the scratch as compared to cells 100µm from the edge of the scratch after one hour of migration (Figure 6–figure supplement 1D). MCF7 cells are roughly 20-25µm in diameter, thus the elevated RhoA near the scratch edge likely represents leader cells. Depletion of *VANGL2* significantly reduced RhoA activity in leader cells 5µm from the edge of the scratch (Figure 6C) and appeared to reduce RhoA activity in follower cells 100 µm from the edge of the scratch, but did not pass our threshold for statistical significance (Figure 6D) after one hour of migration. Rac1 signaling did not differ spatially within the migrating cohort and modulation of *VANGL2* did not alter Rac1 activity (Figure 6–figure supplement 1A). Collectivly, these findings suggest that Vangl2-dependent Wnt/PCP signaling specifically regulates RhoA in leader cells to support actin cytoskeletal rearrangements critical to the formation of pro-migratory protrusions that drive collective migration.

Mechanistically, our observations demonstrate that Vangl2 acts at the leading edge of collectively migrating cells to prompt the RhoA-dependent cytoskeletal dynamics necessary for pro-migratory cellular protrusion formation (Figure 7). By extension, we propose that the high Vangl expression frequently observed in primary tumors drives aberrant Vangl-dependent Wnt/PCP signaling, resulting in a hyper-protrusive leading-edge that supports invasiveness and ultimate metastatic dissemination.

**Figure 7.**
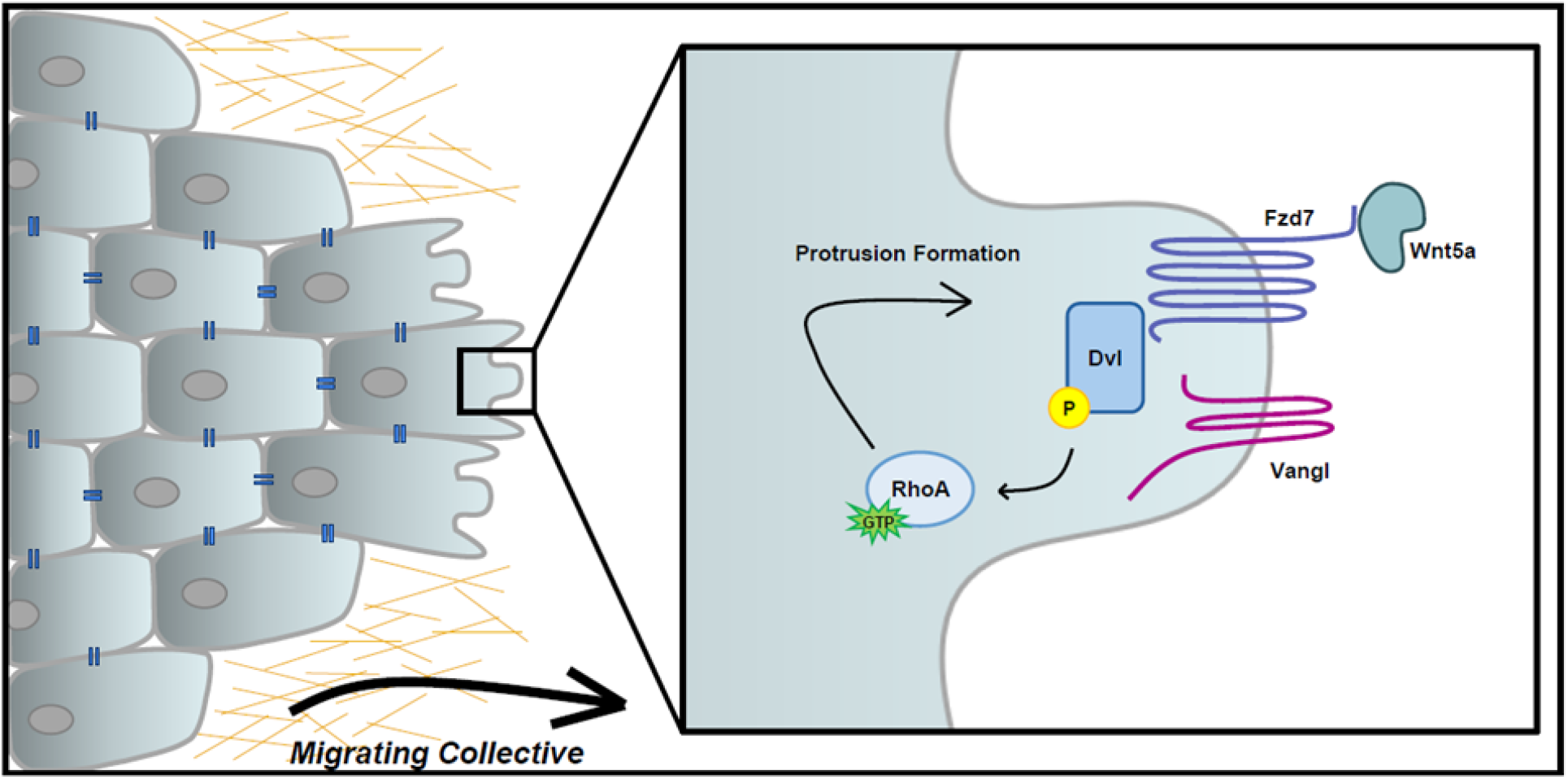
Model of Vangl-dependent Wnt/PCP signaling mediating breast cancer cell collective migration and invasion. Vangl2 acts at the leading edge of collectively migrating cells to prompt the RhoA-dependent cytoskeletal dynamics necessary for pro-migratory cellular protrusion formation.

## Discussion

Metastatic disease is responsible for the majority of cancer-related deaths (Steeg, 2016), despite significant investment into elucidating molecular drivers of metastasis and identifying opportunities for therapeutic intervention. The acquisition of migratory and invasive behaviors by tumor epithelial cells is a critical step for metastatic dissemination, but the molecular mechanisms underlying this transition remain incompletely described. However, tumor cells commonly reactivate and exploit developmental motility programs to promote malignancy, giving rise to invasiveness, metastasis, and poor patient survival (Friedl & Gilmour, 2009). In this study, we demonstrate that Wnt/PCP signaling, which in embryos promotes cell polarity and motility to regulate proper tissue structuring (Devenport, 2014), drives the collective migration and invasion of breast tumor cells. We show that the Wnt/PCP-specific transmembrane scaffold protein Vangl2 is vital for the efficient formation of metastatic lung lesions from ErbB2-driven mammary tumors, and localizes to the leading-edge of cohorts of collectively migrating breast cancer cells where it engages RhoA to drive the formation of pro-migratory protrusions.

Despite reports that Vangl2 is highly expressed in 25% of invasive breast cancers (Hatakeyama et al., 2014) and that elevated *VANGL2* correlates with advanced stage disease and decreased metastasis-free survival of breast cancer patients (Puvirajesinghe et al., 2016), the functional role of Vangl2 in breast cancer malignancy has remained largely unexplored. Here, we present the novel finding that that Vangl2, a critical component of Wnt/PCP signaling, is vital to efficient metastasis of mouse mammary tumors. Loss of *Vangl2* does not impact the efficiency with which disseminated tumor cells colonize the lung nor the proliferation of established metastatic lesions. We instead find that high Vangl expression drives collective cell migration *in vitro* and collective cell invasion *ex vivo,* and elevated *Vangl1* and *Vangl2* expression in K14-positive invasive leader cells *in vivo*. High Vangl expression results in hyper-protrusive leading-edge morphology within leader cells of collectively migrating cohorts suggesting that Wnt/PCP signaling is critical in promoting the cytoskeletal remodeling necessary for the invasive behavior of primary breast tumor cells. Our data indicate that Vangl2 contributes to local collective and migration within the primary tumor, consistent with its role in mediating cell motility during embryonic development (Butler & Wallingford, 2017), resulting in increased dissemination from the primary tumor.

While *Vangl2* is critical for metastasis of *MMTV-NDL* tumors, it is dispensable for both the initiation and growth of the primary tumor. Our observations are consistent with a role for Vangl2 in mediating cell motility events critical to developing embryos (Butler & Wallingford, 2017; Devenport, 2014), but conflict with a previous report in which shRNA-mediated knockdown of *VANGL2* decreased proliferation of SUM159 and HCC1806 breast cancer cells xenografted into the flanks of NSG mice (Puvirajesinghe et al., 2016). Based on reports that *VANGL2* upregulation is associated with higher grade breast tumors (Puvirajesinghe et al., 2016), we hypothesize that elevated *VANGL2* expression and resulting activation of Wnt/PCP signaling is a feature of late-stage or advanced disease. Because the autochthonous *MMTV-NDL* model recapitulates the full course of breast cancer development beginning from an untransformed mammary epithelial cell, Wnt/PCP signaling may have not yet been aberrantly engaged for the majority of primary tumor growth. Xenograft tumors derived from cell lines models of late-stage breast cancer may more heavily rely upon Wnt/PCP signaling for primary tumor growth. This discrepancy suggests that Wnt/PCP signaling may be a marker that could be used clinically to predict invasive or aggressive breast cancer.

Our finding that elevated Vangl protein mediates phosphorylation of Dvl2, a critical effector of downstream Wnt/PCP signaling, led us to speculate that Vangl may directly regulate actin cytoskeleton rearrangements in leader cells. Both the Rho GTPases Rac1 and RhoA are essential components of Wnt/PCP signaling-mediated motility during embryonic development (Habas et al., 2003). To investigate Rho GTPase spatiotemporal dynamics during breast cancer collective cell migration, we employed time-lapse microscopy of MCF7 cells stably expressing FRET biosensors for Rac1 and RhoA. We measured the spatial activity of each GTPase as a function of distance from the edge of a scratch in the cellular monolayer throughout a 12-hour scratch migration. We found RhoA activity consistently peaks in leader cells, whereas Rac1 has a more uniform spatial activity pattern across both leader and follower cell populations. Observation of elevated RhoA activity in MCF7 leader cells was somewhat surprising, as leader cells in collective cell migration are generally thought to have high Rac activity at the cell front to coordinate membrane protrusions via branched actin polymerization, while high RhoA activity at the cell rear facilitates acto-myosin based contraction (Karlsson et al., 2009; Mayor & Etienne-Manneville, 2016; Zegers & Friedl, 2014). However, spatial patterning of Rac1 and RhoA activity appear to be specific for both cell migration mode and cell type. For example, in individual migrating normal kidney epithelial Ptk1 cells, RhoA plays a central role in driving anterograde actin flows and contractile forces on strong focal adhesions at the cell front, which work to pull the cell forward (Gupton & Waterman-Storer, 2006). In collectively migrating Madin-Darby Canine Kidney (MDCK) cells, thick actin-myosin II cables around the perimeter of protrusive cell collections termed ‘fingers’ and RhoA activity highest in leading-edge cells regulating the acto-myosin cable (Reffay et al., 2014). Future work is required to determine if collectively migrating MCF7 cells, or collectively migrating breast cancer cells more generally, develop acto-myosin cables analogous to MCDK cells.

Our findings provide substantive insight into the mechanism by which Vangl proteins specifically mediate the formation of pro-migratory protrusions at the leading-edge of leader cells in collectively migrating cohorts. Additionally, these data suggest that Vangl-mediated regulation of RhoA dynamics in leader cells is critical to Wnt/PCP-mediated collective cell migration and invasion. However, the molecular underpinnings of Vangl-mediated RhoA activity within leader cells are not clear. In vertebrate gastrulation, Wnt/PCP signaling appears to drive cellular motility via engagement of Rho family GTPases in a manner that depends on both the cytoplasmic effector Dvl and Daam1, a Formin homology protein that mediates Wnt-induced Dvl-Rho complexes (Habas et al., 2001). While this study did not investigate whether Vangl is a required component of this complex, we speculate that Vangl may serve as a required scaffold upon which the Dvl-Daam1-RhoA complex assembles in leader cells. This is consistent with previous reports suggesting that Vangl may be a master scaffold upon which diverse complexes assemble (Anastas et al., 2012; Puvirajesinghe et al., 2016; Wald et al., 2017). These findings, coupled with our observations that Vangl localizes to leader cells in collectively migrating and invading cohorts and drives the formation of pro-migratory protrusions, suggests that Vangl-RhoA-mediated modulation of the cytoskeleton in leader cells is a significant contributor to the invasive nature and metastatic dissemination of primary tumor cells.

## Materials & Methods

### Generation of Vangl2/NDL mice

All experimental protocols were approved by the IACUC of the University of California, Davis, CA, USA. *Vangl2^tm2.1Mdea^/J* conditional knockout mice (Copley et al., 2013) (The Jackson Lab, Stock #025174) were crossed with *Tg(MMTV-cre)4Mam/J* mice (Wagner et al., 1997) (The Jackson Lab, Stock #003553) to generate mice with Vangl2 deletion in the mammary gland. The *MMTV-NDL* mouse has been previously described (Siegel et al., 1999). Genotypes were confirmed by polymerase chain reaction in house using primers for Vangl2 (Fwd 5’-CAGAA CCTCCTGTCCCTGA-3’; Rev 5’-CTCAGCTAAACCACCTCTGC-3’), Cre (Fwd 5’-GCGGTCTGGCAGTAAAAACTATC-3’; Rev 5’-GTGAAACAGCATTGCTGTCACTT-3’), and NDL (Fwd 5’-TTCCGGAACCCACATCAG -3’; Rev 5’- GTTTCCTGCAGCAGCCTA -3’).

### Tumor monitoring and analysis

Mammary tumors were palpated once or twice weekly in female Vangl2/NDL mice commencing at 16 weeks of age by a single investigator and all palpable tumors were measured by calipers. When the largest tumor reached 2cm in any direction, mice were euthanized by CO_2_ asphyxiation and tumors were collected and either fixed in 10% neutral buffered formalin for paraffin embedding and sectioning or further dissociated for *in vitro* analysis. Mice with illnesses arising independent of their tumors that required sacrifice prior to reaching the pre-determined endpoint were excluded from analyses.

### Histology and Immunohistochemistry

Histologic analysis of lungs was performed for all mice on study (n=20 per genotype) and for a randomly selected subset of Vangl2^+/+^/NDL and Vangl2^fl/fl^/NDL mammary tumors (n=4 per genotype). H&E-stained sections were prepared using previously described methods (Rowson-Hodel et al., 2015). Immunohistochemistry (IHC) was performed as previously described (Rowson-Hodel et al., 2018). An internal negative control (no primary antibody) was included with each assay.

### Lung metastases analysis

Lungs were inflated with PBS, fixed in 10% neutral buffered formalin, paraffin embedded, and sectioned for IHC analysis and H&E staining. The number of ErbB2- positive metastatic lesions present on all five lung lobes were counted for all mice on study (n=20 per genotype) from images taken on a Keyence BZ-X810 microscope. Metastatic burden was quantified by normalizing the number of metastatic lesions to the total tumor burden.

### Tail vein injections

Pooled primary Vangl2^+/+^/NDL and Vangl2^fl/fl^/NDL mammary tumors (Vangl2^+/+^/NDL *n*=11, Vangl2^fl/fl^/NDL *n*=10; 5 x 10^5^ in 200 µL PBS) were instilled to the lateral tail vein of 12-week-old FvB/NJ mice. Lungs were harvested 6-weeks post injection and analyzed for metastatic lesions. Mice were randomly assigned to cohorts and were caged as mixed cohorts.

### Cell culture and reagents

BT549, HCC1937, HCC38, MCF7, MDA-MB-157, MDA-MB-231, MDA-MB-468, nMuMG, L-Cells, and L-Cells-Wnt3a, and HEK293T cells were purchased from the American Type Culture Collection (ATCC) and maintained as recommended at 37°C in 10% CO_2_ in media supplemented with 10% fetal bovine serum (FBS, Genesee Scientific), 1% penicillin-streptomycin (Invitrogen). Met-1 (gifted by A.D. Borowsky) and NDL cells were maintained as previously described(Borowsky et al., 2005; Miller et al., 2008). Prior to use, cell lines were authenticated by short-tandem repeat profiling (Genetics Core Facility; University of Arizona, Tucson, AZ, USA) and tested for mycoplasma contamination by RT-PCR as described (Uphoff & Drexler, 2002, 2004). Antibodies used for immunoblotting, immunofluorescence, and immunohistochemistry are as follows: anti-Dvl2, anti-p-β-Catenin (Ser33/37/Thr41), anti-β- Catenin (Cell Signaling), anti-Tubulin (Sigma), horseradish peroxidase-conjugated goat anti-mouse and goat anti-rabbit secondary antibodies (Bio-Rad), anti-Vangl1 (R&D Systems), anti- Fzd7 (Abcam), anti-Keratin14 (Biolegend), anti-E-Cadherin, anti-Flag and anti-V5 (Cell Signaling), anti-Phalloidin647 (Invitrogen), and AlexaFluor 488-conjugated goat anti-mouse, AlexaFluor 546-conjugated goat anti-rabbit, and Alexa-Fluor 568-conjugated goat anti-chicken secondary antibodies (Invitrogen), and anti-Ki67, anti-c-Caspase3, and anti-ErbB2 (Cell Signaling). Cells were treated with the Wnt-inhibitor C59 at 100nM (R&D Systems).

### Generation of stable overexpression and knockdown cell lines by lentiviral transduction

*VANGL2*-targeted shRNA constructs shVangl2-1 (ID: V3LHS_334647 or ID: TRCN0000180101), shVangl2-2 (ID: V3LHS_334648 or TRCN0000417141) (Dharmacon, Sigma-Aldrich), or control vectors pGIPZ (Dharmacon) or pLKO.1 (a gift from David Sabatini, Addgene plasmid #1864; http://n2t.net/addgene:1864; RRID:Addgene_1864) were employed for Vangl2-depletion studies. Stable overexpression cells were created using Vangl1 and Vangl2 plasmids (Harvard PlasmID repository, HsCD00339551 and HsCD00294893) and Wnt5a plasmid that was a gift from Marian Waterman (Addgene plasmid # 35911; http://n2t.net/addgene:35911; RRID:Addgene_35911) subcloned into the control vector pLX304, a gift from David Root (Addgene plasmid # 25890; http://n2t.net/addgene:25890; RRID:Addgene_25890). VSVG-pseudotyped lentivirus was generated by transfecting HEK293T cells with the psPax2 packaging vector. Cells were transduced with 10µg/mL polybrene (Millipore), followed by drug selection with 1µg/mL Puromycin (Sigma-Aldrich) or 4µg/mL Blasticidin (Sigma-Aldrich).

### Wnt5a and Wnt3a stimulation

Wnt5a conditioned media was produced by stably transducing nMUMG, MCF7, MDA-MB-231, MDA-MB-468, or NDL cells with Vector- or Wnt5a-containing lentivirus. Vector- or Wnt3a- conditioned media was collected from L-Cell and L-Cell-Wnt3a, respectively. Conditioned media was collected from confluent cell culture plates, cleared of debris by centrifugation, and stored at -80°C.

### Scratch migration assays

Confluent monolayers of cells were scratched with a sterile pipette tip and imaged immediately and after twelve hours with an Olympus IX81 microscope with CellSens Entry software. Scratch area was measured using ImageJ (NIH) and the area of the scratch filled in over twelve hours was quantified. Results were normalized to appropriate controls for each assay.

### Immunoblotting

Cells were washed with 1X PBS and lysed directly in 2x Laemmli sample buffer. All samples were resolved by SDS-PAGE, transferred to nitrocellulose membranes, and blotted with the indicated antibodies. Immunoblots were developed using Pierce SuperSignal West chemicals (Thermo Fisher) on an AlphaInnotech imaging station and quantified with ImageJ (NIH).

### Real-time PCR analysis

RNA was collected using a PureLink RNA MiniKit (Ambion) and converted to cDNA with the High-Capacity cDNA reverse transcription kit (Applied Biosystems). *q*PCR was conducted in a Bio-Rad CFX96 real-time PCR system using TaqMan gene-specific primer/probe sets (Applied Biosystems) and SsoAdvanced master mix (Bio-Rad). Analysis was conducted using Bio-Rad CFX Manager software and message levels were normalized to GAPDH.

### *Ex vivo* 3D organoid invasion assays

*MMTV-PyMT* tumor samples were a kind gift from Dr. Jason Hatakeyama (Stanford University, Stanford, CA, USA). Tumors were dissociated into single cells as previously described(Diehn et al., 2009) with minor modifications and seeded in Matrigel (Corning) with organoid growth media, which has been previously described (Nguyen-Ngoc et al., 2012). After twenty-four hours in Matrigel, cells were transduced with specified lentivirus and spinfected for one hour at ∼500G in a Beckman centrifuge. After seven days in culture, organoids derived from single *MMTV-PyMT* tumor cells were recovered from Matrigel using Cell Recovery Solution (Corning) and embedded into rat-tail collagen I (Thermo Fisher) as previously described (Nguyen-Ngoc et al., 2012). Invasive protrusions were imaged and counted with an Olympus IX81 microscope with CellSens Entry software or Zeiss LSM 710 AxioObserver confocal microscope.

### RNA-seq data mining

Raw RNAseq reads from Cheung et al., archived as SRP066316 was downloaded from the Sequence Read Archive (Cheung et al., 2016). Reference genome for pseudoalignment was built in Kallisto v0.43.1 from Genome Reference Consortium Mouse Build 38 using a k-mer length of 31. Reads were then pseudoaligned to the reference genome using 100 bootstraps to estimate error. Differential expression analysis was then performed in R using the DESeq2 package (1.28.1). Biological replicate #3 (SRR291722 and SRR2921727) varied considerably from the other replicates by principal component analysis and expression of key marker genes, and was therefore omitted from the analysis.

### Immunofluorescence microscopy

Cells were seeded onto coverslips, fixed with 4% paraformaldehyde, and stained as indicated. PyMT-derived organoids embedded in collagen I were fixed with 4% PFA and stained as indicated. Imaging was conducted on a Zeiss LSM 710 AxioObserver confocal microscope or Keyence BZ-X810 microscope. An internal negative control (no primary antibody) was included with each assay. Average number of Vangl1-rich protrusions/cell was quantified by counting the number of Vangl1+ protrusions in leader cells along the leading-edge of a MCF7 cells actively migrating into a scratch made in a confluent monolayer from 4 or 8 independent scratch assays. Percentage of cells with Vangl1-rich protrusions was quantified by counting the number of leader cells with and without Vangl1+ protrusions along the leading-edge of MCF7 cells actively migrating into a scratch made in a confluent monolayer from 4 or 8 independent scratch assays.

### FRET biosensor scratch assay imaging

Rac1 and RhoA intramolecular FRET biosensors and have been previously described (Itoh et al., 2002). MCF7 cells stably expressing *VANGL2-* targeted shRNAs to deplete Vangl2 were stably transfected with Rac1 or RhoA intramolecular FRET biosensors using PEI transfection reagent. Rac1 or RhoA biosensor expressing cells were sorted with a BD “inFlux” 18-color cell sorter (Becton Dicksinson). For all imaging experiments, cells were plated on glass-bottomed 96-well plates (Cellvis) and grown to confluency. Prior to imaging, the monolayers were scratched with a sterile pipette tip, and the media was replaced with Liebovitz-15 (L-15) media, which was made with no riboflavin, folic acid or dyes to reduce autofluorescence from the media supplemented with 2% FBS (UC Davis Biological Media Services). The plates were then transferred to a Nikon Eclipse TI equipped with an OKO Labs cage incubator set to 37 °C. The microscope is controlled by MATLAB (version 2015 A) through Micro-Manager (v 1.6), allowing precise, repeatable experiments. The X & Y stage positions of the scratch were identified by the user and all scratch positions were stored in MATLAB for time lapse imaging. Epifluorescent CFP/YFP FRET images were collected every 15 min for 12 hours using a 20x Nikon Apochromat 0.75 NA objective. Cyan (∼440 nm) excitation illumination was provided by the X-Cite XLED1 BLX module, while simultaneous acquisition of FRET images was achieved using dual Andor Zyla 4.2 sCMOS cameras separated by a Cairn TwinCam LS image splitter with a Chroma Technologies dichroic mirror (ZT491rdc) that reflects wavelengths less than 502nm to one camera, while passing longer wavelengths to the second camera.

### Camera and illumination corrections

The dark-state noise for each camera was empirically measured as described (Bell et al., 2021). In brief, several images were captured without illumination and the microscope light path set to the oculars. The dark-state correction image was generated by taking the median over the stack of dark images. This correction was then subtracted from all experimental images. CFP/YFP FRET ratio images were observed to have a gradient of activity from the top to the bottom of the images. A correction image was developed to remove this gradient as described (Bell et al., 2021). Images of unstimulated, confluent monolayers of MCF7 cells expressing the CFP/YFP FRET sensor were collected. FRET ratio images were generated from raw CFP and YFP images that were processed using our standard analysis pipeline. The median FRET ratio was taken over the stack of images on a pixel-by-pixel basis. Only pixels that overlapped with a cell logical mask were included in the median analysis. To reduce local variability effects and noise, the median image was broken into 24×24 pixel blocks. Next, the median was taken for each block, the resulting image was then smoothed using a gaussian filter (sigma=5) and the image was then resized to match the size of the input image. To apply the correction, experimental FRET ratio images were divided by the ratio correction image.

### Image alignment, background subtraction, segmentation, and speckle filtering

All image analysis methods were conducted using MATLAB. CFP/YFP FRET images were aligned using an alignment algorithm as described (Yang et al., 2016). Images were then cropped to ensure equal size. To estimate the background, empty wells containing L-15 media + 2% FBS were imaged with CFP/YFP imaging configurations that were identical to the experimental conditions. Eight empty well frames were collected, and the median was calculated on a per pixel basis over the image stack. The median well background images were then aligned and cropped to match the size of the experimental images. Next, the background mask was determined for the experimental images. To generate the background mask, the experimental images were log transformed to enhance the dimmer cell pixels. The image threshold was then calculated using Otsu’s method (Otsu, 1979). The background logical mask was created by finding pixels in the log transformed image below the threshold. Pixels contained in the background mask were used to find the median intensity in both the empty well image and the experimental image. The ratio of the two median intensities was then used to scale the empty well image to match the intensity of the background pixels in the experimental image. Once scaled, the empty well images were subtracted from the experimental images to remove background.

To segment the cells for further processing, the scaled background image was subtracted from the CFP and YFP images. The CFP and YFP images were added to reduce signal to noise, and the sum image was then used to identify the cell mask and the scratch mask. Because the cell pixels were much brighter than the scratch pixels, the dimmest 0.5% of pixels were subtracted from the sum image, and the minimum pixel intensity value was set to 20 prior to log transformation. The log transformed images were rescaled to a pixel range from the first percentile to the ninetieth percentile and the background vs foreground threshold was identified using Otsu’s method. Background pixels were identified below the threshold to create a background logical mask. The background mask was morphologically closed to remove small gaps in the mask. Small objects below 50 pixels in area were removed from the mask and small holes in the mask were filled. The inverse of this mask was used to define the cell mask. Both cell and background masks were saved for additional processing.

The raw CFP and YFP images from MCF7 cells had small, but very bright puncta. A speckle filter was developed to remove these puncta from the processed CFP and YFP images. The sum image of the two FRET channels was filtered using a Laplacian operator (alpha=0.9) to convert the brightest pixels to the smallest negative pixels. The Laplace filtered image was subtracted from the sum image, effectively making the brightest pixels even brighter. To threshold rare, but bright pixels, the histogram of the image was taken and the intensity value for the first bin with 9 or fewer pixels with positive intensity values were used as the threshold. A bright pixel binary mask was created for all pixels above the bright pixel threshold in the sum image. The bright pixel mask was then morphologically closed to connect neighboring pixels, and objects greater than 300 pixels were removed. Finally, the FRET reporters are plasma membrane associated, and in some circumstances, were bright enough to be captured by the bright pixel threshold. These objects were more linear as they were essentially tracings of cell edges. To remove these cell membrane objects, the eccentricity for each object was measured. Objects with an eccentricity >0.6 (more linear than circular) were removed from the mask. The speckle mask was then saved for processing the FRET ratio images.

Finally, to create the FRET ratio images, raw CFP and YFP images were background subtracted using the scaled background image constructed above. For both CFP and YFP images, pixels in the background mask and pixels in the speckle mask were set to NaNs to remove them from further processing. The background subtracted and segmented CFP and YFP images were smoothed with a gaussian filter (sigma=2) and saved for further processing. Next, the FRET ratio was calculated by dividing the YFP image by the CFP imaged. The ratio correction image described above was then applied to the FRET ratio image. These FRET ratio images were written to movies and exported for later use.

### Computation of FRET ratios as a function of distance from the scratch edge

To understand whether cells near the scratch had higher GTPase activity when compared to cells in the monolayer, FRET ratios were measured and binned based on their distance from the scratch edge. To identify the scratch edge, the background masks described above were further analyzed. The scratch is the largest object in the background mask, thus the object with the maximum area was defined as the scratch mask. Next, all other objects in the background mask were removed and the perimeter from the scratch mask was defined. The scratches were consistently generated North to South on the well, thus pixels touching the top and bottom of the perimeter mask image were removed, leaving two scratch edge perimeter masks that corresponded with the left and right sides of the scratch. The scratch edges were then dilated by 2 pixels to ensure overlap with the edge of the cell mask defined above. Additionally, the cell masks were separated based on their relative position to the scratch mask (Left or Right). The MATLAB bwdistgeodesic function was then used to measure the “quasi-euclidean” distance of all pixels in the left and right cell masks based on their distance from the corresponding scratch edge mask. These distance masks were used to sort pixels into 5 µm bins based on their distance from the scratch edge. The ratio correction image was applied to the YFP images, then the mean intensity values for the background subtracted, and segmented FRET donor and FRET acceptor images were calculated for each bin. The mean FRET ratio for each bin was then calculated. These measurements were compiled for each well within the experimental groups and were used to generate the plots reported.

## Statistical analysis

Prism software (GraphPad Software) was used for all statistical analyses. Statistical significance was determined by two-sided unpaired t-test with Welch’s correction, paired t-test, Mann-Whitney test, Log-rank test, or likelihood ratio test followed by Benjamin-Hochberg correction for multiple hypothesis testing (indicated in the figure legends). *P*-values ≤ 0.05 were considered statistically significant.

## Acknowledgements

These studies were supported by NIH grants R01CA230742 (KLC), F31CA210467 (KV), F31CA246900 (CD), F31CA165758 (JH), T32GM099608 (AB), and DP2HD094656 (SRC) and NSF Graduate Research Fellowship 1650042 (GRRB). We thank Henry Ho for discussion, reagents, and critical reading of the manuscript. We are grateful to Jonathan Van Dyke and the UCD Comprehensive Cancer Center Flow Cytometry Shared Resources supported in part by NIH grant P30CA093373 and Gabe Jackman at Keyence for their technical assistance.

## Author Contributions

K.V., J.H., and K.L.C. contributed to conceptualization; K.V., J.H., G.R.R.B., S.R.C., and K.L.C. contributed to methodology; K.V., C.A.D., G.R.R.B., A.B., M.H., and H.L. contributed to data collection; K.V., C.A.D., J.H., G.R.R.B., S.R.C., and K.L.C. contributed to data analysis and interpretation; S.R.C. and K.L.C. contributed resources; K.V., J.H., G.R.R.B., and K.L.C. contributed to manuscript writing.

## Competing Interests

The authors declare no competing interests.

**Supplementary Video 1. RhoA-FRET biosensor video of MCF7-Control.** Representative videos of spatial activity profiles of RhoA in collectively migrating MCF7 cells stably expressing RhoA-FRET biosensor for Control over 12 hours. Color bars indicate the range of RhoA-FRET biosensor ratios. Scale bar=25µm.

**Supplementary Video 2. RhoA-FRET biosensor video of MCF7-shVangl2-1.** Representative videos of spatial activity profiles of RhoA in collectively migrating MCF7 cells stably expressing RhoA-FRET biosensor for shVangl2-1 over 12 hours. Color bars indicate the range of RhoA- FRET biosensor ratios. Scale bar=25µm.

**Supplementary Video 3. RhoA-FRET biosensor video of MCF7-shVangl2-2.** Representative videos of spatial activity profiles of RhoA in collectively migrating MCF7 cells stably expressing RhoA-FRET biosensor for shVangl2-2 over 12 hours. Color bars indicate the range of RhoA- FRET biosensor ratios. Scale bar=25µm.

**Immunoblot Source Files.** The raw data corresponding to Figures 2D, 3E and 3G, and Figure 2-figure supplement 1C, Figure 3-figure supplements 1A and 1B, and Figure 4-figure supplements 1A and 1B, are compiled in eight folders. Regions of blots included in figures are indicated by red boxes in two Word documents.

---

**Figure 1–figure supplement 1.**
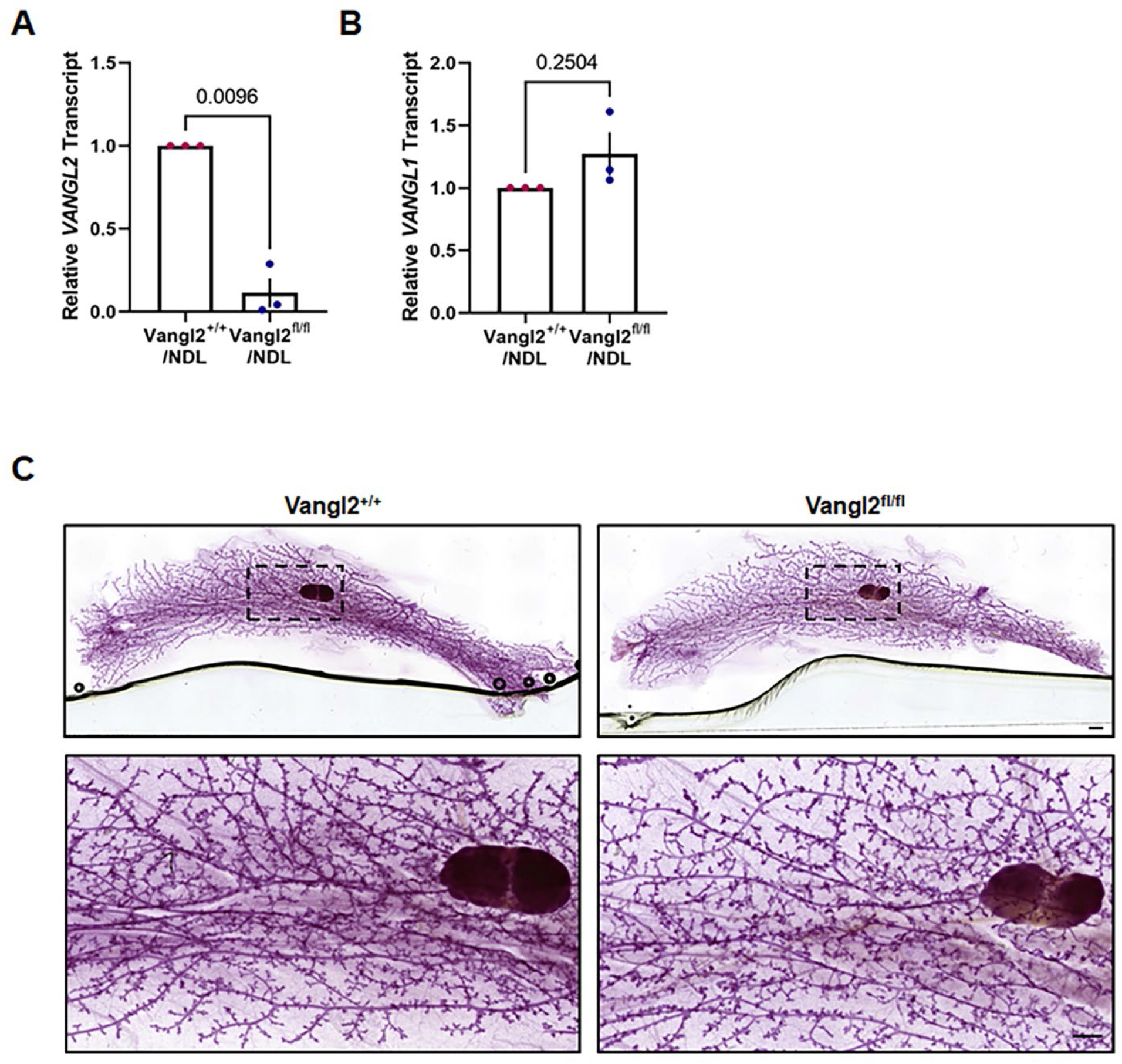
Functional Vangl2 deletion is evident by transcript. **a-b** *VANGL2* **(A)** and *VANGL1* **(B)** transcript from Vangl2^fl/fl^/NDL and Vangl2^+/+^/NDL primary tumors by *q*PCR from three tumors of independent biological sources per genotype. **c** Carmine alum stained mammary whole mounts from estrus matched 20-week-old Vangl2^+/+^ and Vangl2^fl/fl^ mice, demonstrating no detectable differences in gland architecture in adult virgin mice. Scale bars = 1mm (top panels) or 500µm (bottom panels).

**Figure 1–figure supplement 2.**
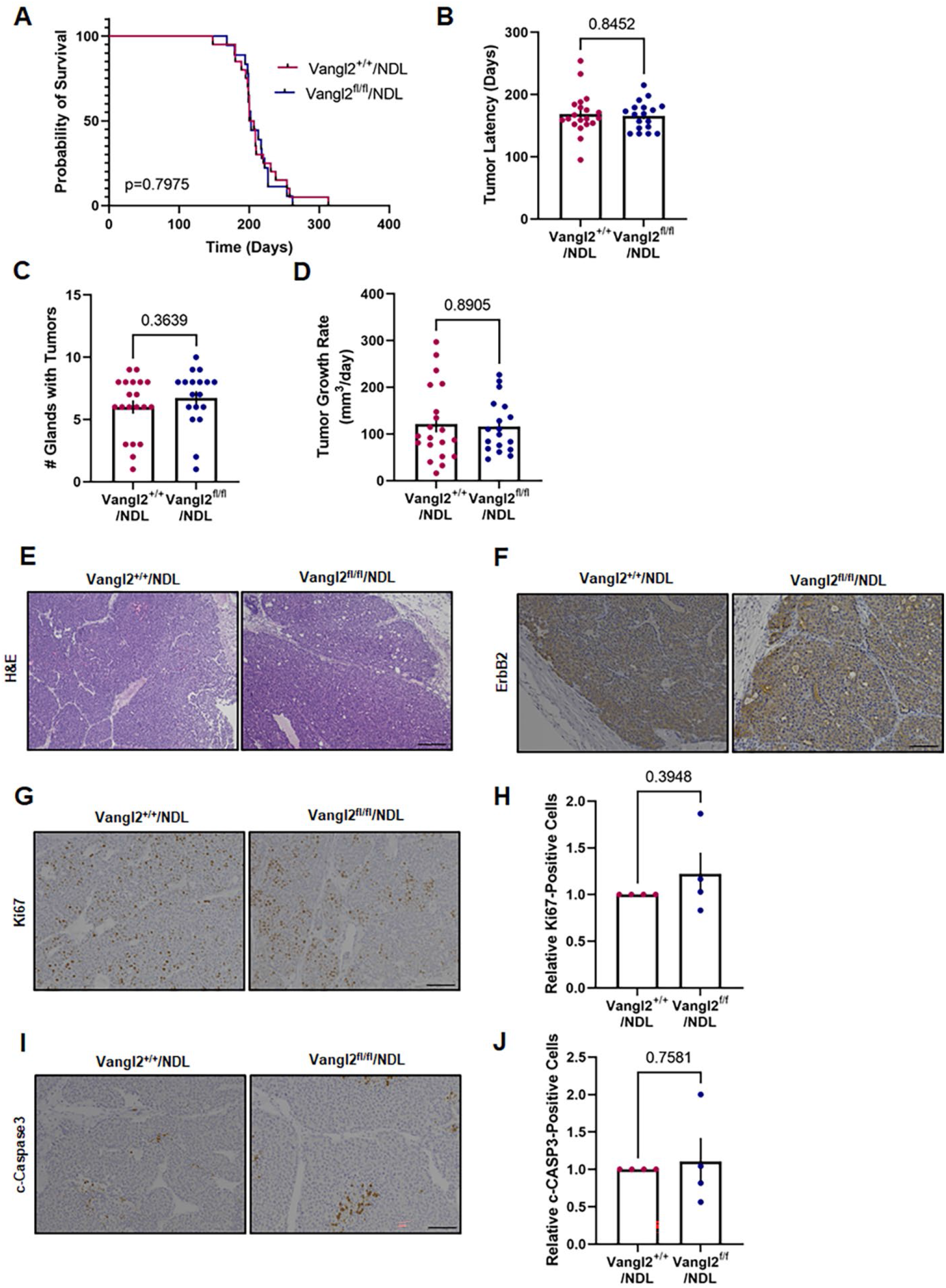
Vangl2 deletion does not impact primary tumor growth characteristics or histology. **A-D** Survival curves and bar graphs depicting Vangl2/NDL tumor initiation and growth characteristics. Probability of survival **(A),** tumor latency **(B),** number of glands with tumors **(C),** and tumor growth rate **(D)** for Vangl2^+/+^/NDL (*n*=20) and Vangl2^fl/fl^/NDL (*n*=20) tumor-bearing animals. **E,F** Representative images are depicted of formalin fixed, paraffin embedded sections from Vangl2^+/+^/NDL and Vangl2^fl/fl^/NDL primary tumors stained with H&E **(E)** or following immunodetection with ErbB2 **(F)**, scale bar=200µm. **G-J** Representative images of Vangl2^+/+^/NDL and Vangl2^fl/fl^/NDL primary tumor tissues following immunodetection of proliferation marker Ki67 **(G)** with quantification of Ki67-positive cells (*n*=4) **(H)** and apoptosis marker cleaved caspase-3 **(I)** with quantification of c-Caspase 3-positive cells (*n*=4) **(J)**, scale bar=100µm. Significance determined by Log-rank **(A)** or Mann-Whitney test **(C,D)** or two-sided unpaired *t*-test with Welch’s correction **(H,J)**. All bar graphs represent the mean ± sem of experimental replicates (*n*).

**Figure 1–figure supplement 3.**
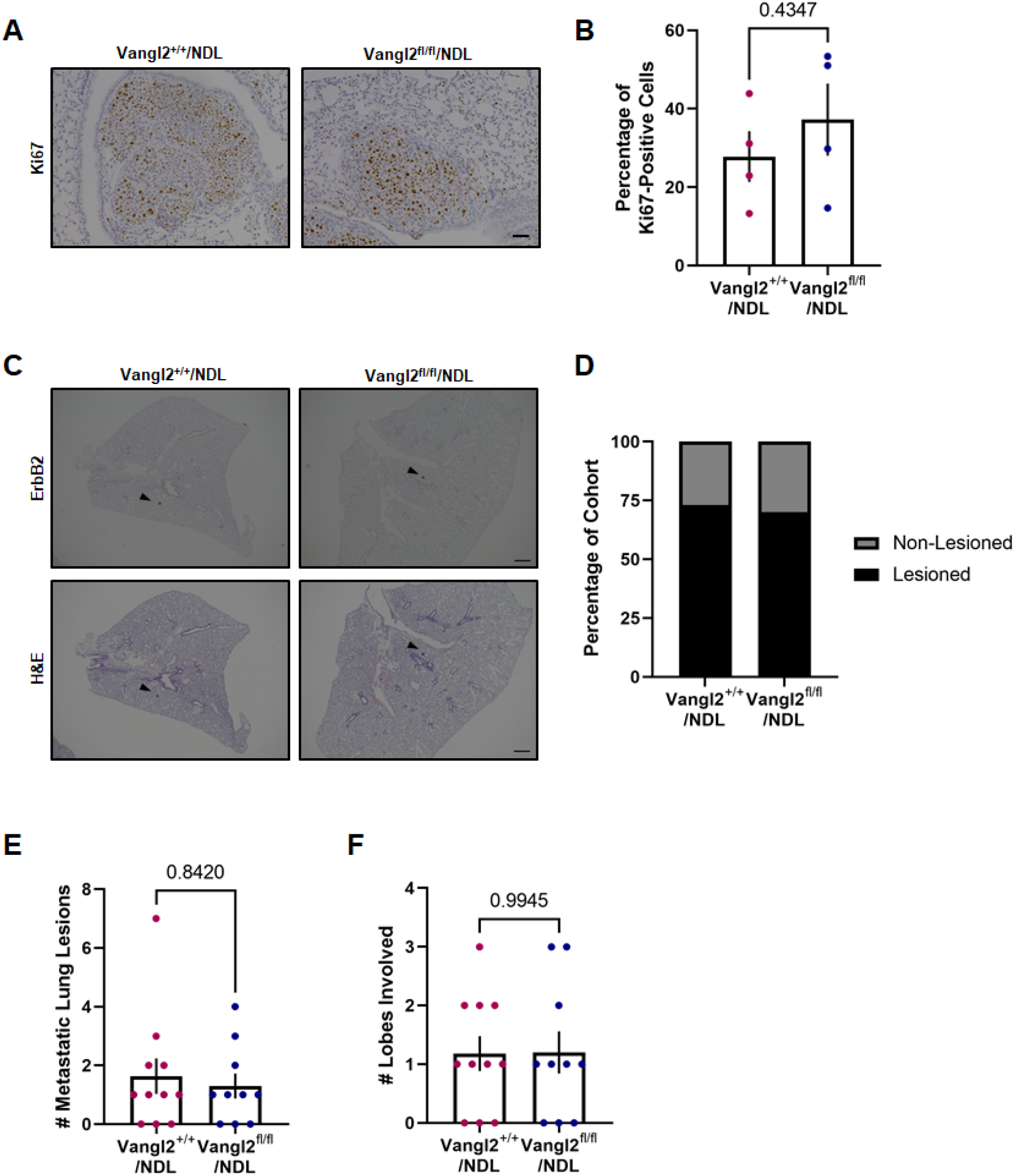
Vangl2 is dispensable for metastatic lesion colonization and proliferation. **A,B** Metastatic lung lesions of Vangl2^+/+^/NDL (*n*=4) and Vangl2^fl/fl^/NDL tumor-bearing mice (*n*=4) were evaluated for proliferative capacity by immunodetection of Ki67 **(A)** and the percentage of Ki67-positive cells was quantified **(B). C** Representative images of formalin fixed, paraffin embedded sections of lungs from FvB/NJ mice receiving Vangl2^+/+^/NDL or Vangl2^fl/fl^/NDL cells via the tail vein following immunodetection of ErbB2 (top panel) and H&E staining (bottom panel). Examples of ErbB2-positive metastatic lung lesions are denoted by black arrowheads, scale bar =500µm. **D-F** Lung lobes (5 lobes per mouse) from FvB/NJ mice receiving Vangl2^+/+^/NDL or Vangl2^fl/fl^/NDL cells via the tail vein were evaluated by histology for the occurrence of metastatic lesions for Vangl2^+/+^/NDL (*n*=11) and Vangl2^fl/fl^/NDL (*n*=10) cohorts. The number of mice bearing metastatic lesions **(D)**, numbers of metastatic lesions **(E)**, and numbers of lung lobes involved **(F)** were assessed. Significance was determined by Mann-Whitney test and bar graphs represent the mean ± sem of experimental replicates (*n*).

**Figure 2–figure supplement 1.**
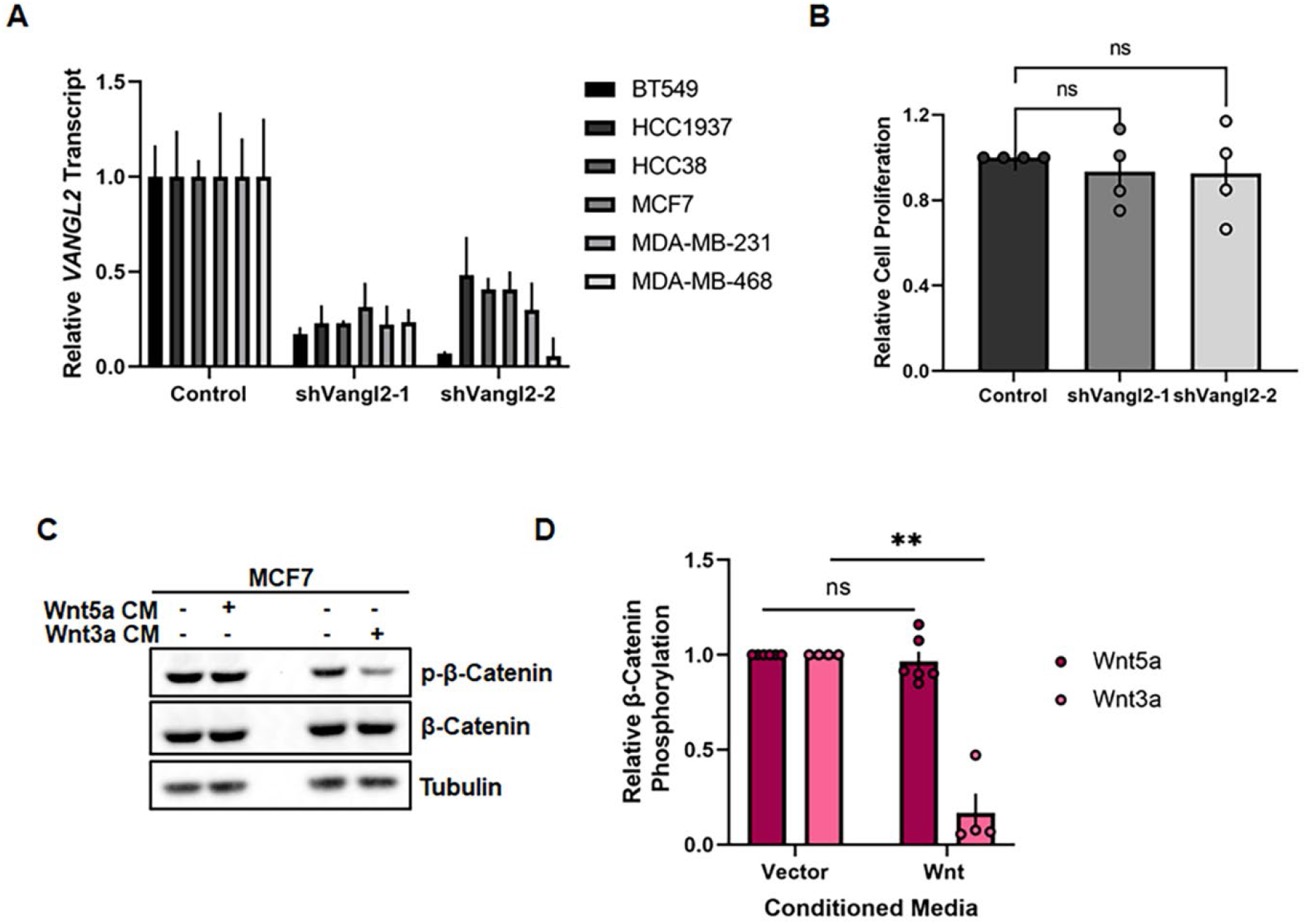
Supporting materials for Figure 2. **A** Relative *VANGL2* transcript in BT549, HCC1937, HCC38, MCF7, MDA-MB-231, and MDA-MB-468 stably expressing Control, shVangl2-1, or shVangl2-2 by *q*-PCR. **B** Quantification of relative cell proliferation of MCF7 cells stably expressing Control, shVangl2-1, or shVangl2-2 (*n*=4). **C,D** MCF7 cells stimulated with Vector- or Wnt5a-conditioned media or Vector- or Wnt3a-conditioned media for 1 hour blotted for β-Catenin and phospho-β-Catenin (Ser33/37/Thr41) **(C)** and quantification of relative phosho-β-Catenin (Ser33/37/Thr41) (control- vs Wnt5a-conditioned media *n*=6, *p*=0.5419, control- vs Wnt3a-conditioned media *n*=4, *p*=0.0038) **(D)**. Bar graphs represent the mean ± sem of experimental replicates (*n*). Significance was determined by a two-sided unpaired *t*-test with Welch’s correction, \**p <* 0.05, \*\**p <* 0.01, \*\*\**p* < 0.001, \*\*\*\**p* < 0.0001.

**Figure 3–figure supplement 1.**
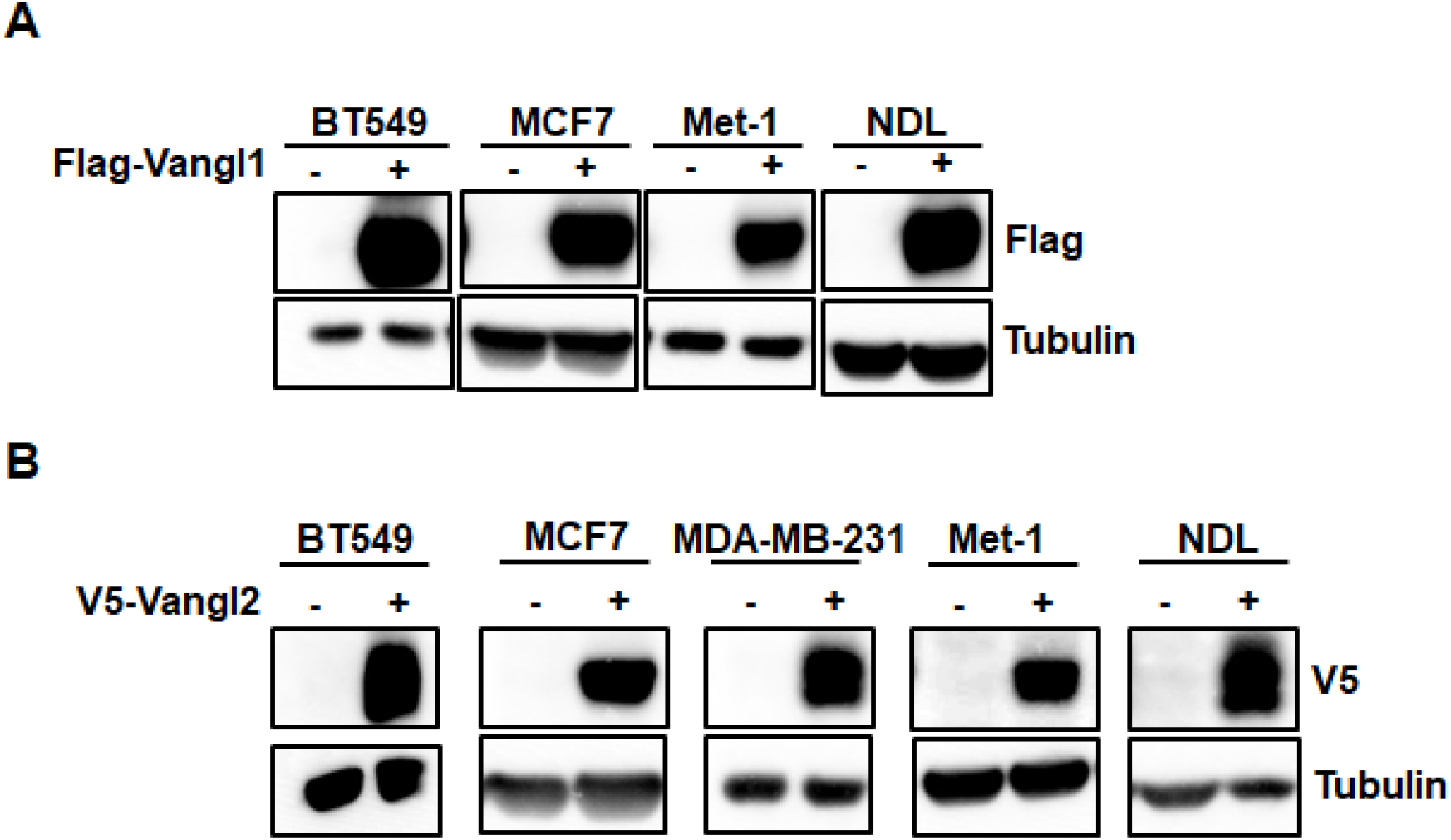
Overexpression of Wnt/PCP components. **A,B** Breast cancer cell lines stably overexpressing Flag-Vangl1 blotted for Flag **(A)** or V5-Vangl2 blotted for V5 **(B).**

**Figure 4–figure supplement 1.**
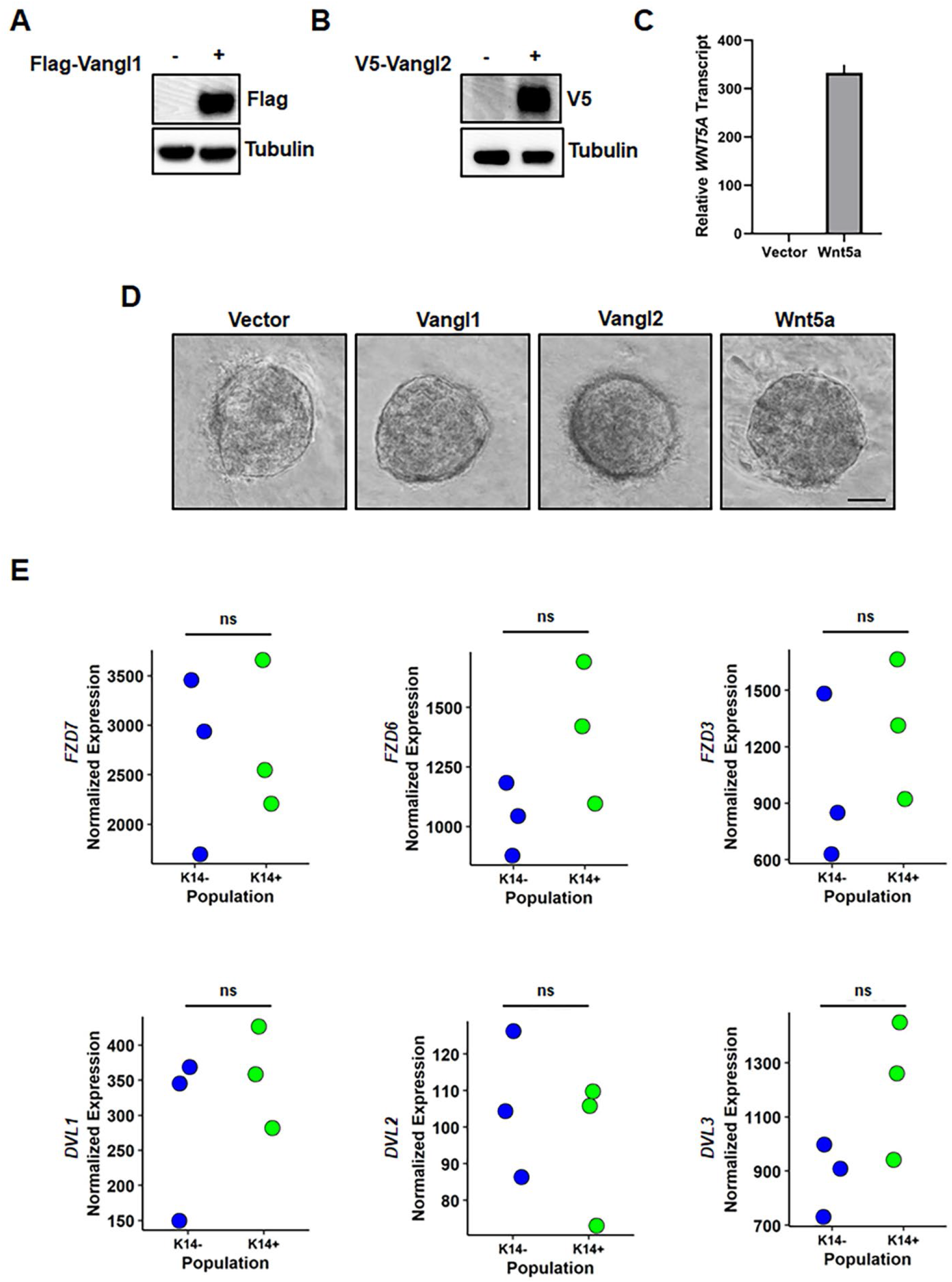
Supporting materials for Figure 4. **A-C** *MMTV-PyMT*-derived tumor organoid cells stably overexpressing Vangl1, Vangl2, or Wnt5a were assessed for Flag- Vangl1 **(A)** or V5-Vangl2 **(B)** expression by Western blot or *WNT5A* transcript by qPCR **(C)**. **D** Representative images of Vector-, Vangl1-, Vangl2-, and Wnt5a-expressing *MMTV-PyMT*- derived tumor organoids in collagen in the absence of *b*FGF, scale bar=50µm. **E** Analysis of RNA-sequencing data set SRP066316 from NCBI Sequence Read Archive for *Fzd7*, *Fzd6, Fzd3, Dvl1, Dvl2*, and *Dvl3* transcript in K14-negative and K14-postive cells derived from *MMTV-PyMT* tumors.

**Figure 5–figure supplement 1.**
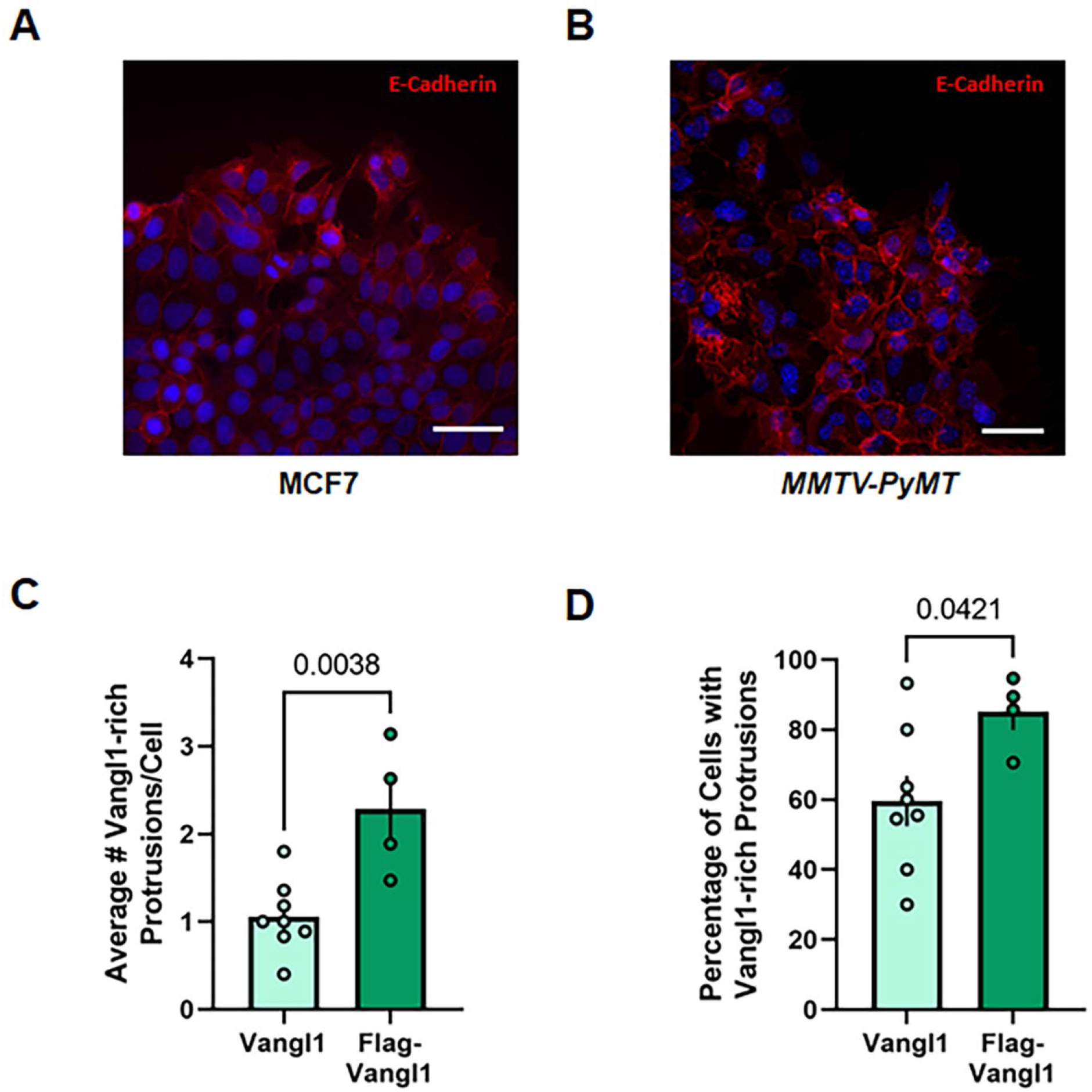
Supporting materials for Figure 5. **A,B** Representative confocal images of collectively migrating MCF7 **(A)** and MMTV-PyMT tumor-derived **(B)** cells stained for E-Cadherin: orange and DAPI: blue, scale bar = 50µm. **C** Quantification of the average number of Vangl1-rich protrusions per leading-edge cell in collectively migrating MCF7-Vector and MCF7-Flag-Vangl1 cells (MCF7-Vector *n*=8 *scratches quantified,* MCF7-Flag-Vangl1 *n*=4 *scratches quantified*, *p=*0.0038) **D** Quantification of the percentage of leading-edge cells with Vangl1-rich protrusions (MCF7-Vector *n*=8, MCF7-Flag-Vangl1 *n*=4, *p*=0.0421). Bar graphs represent the mean ± sem of experimental replicates (*n*). Significance was determined by a two-sided unpaired *t*-test with Welch’s correction.

**Figure 6–figure supplement 1.**
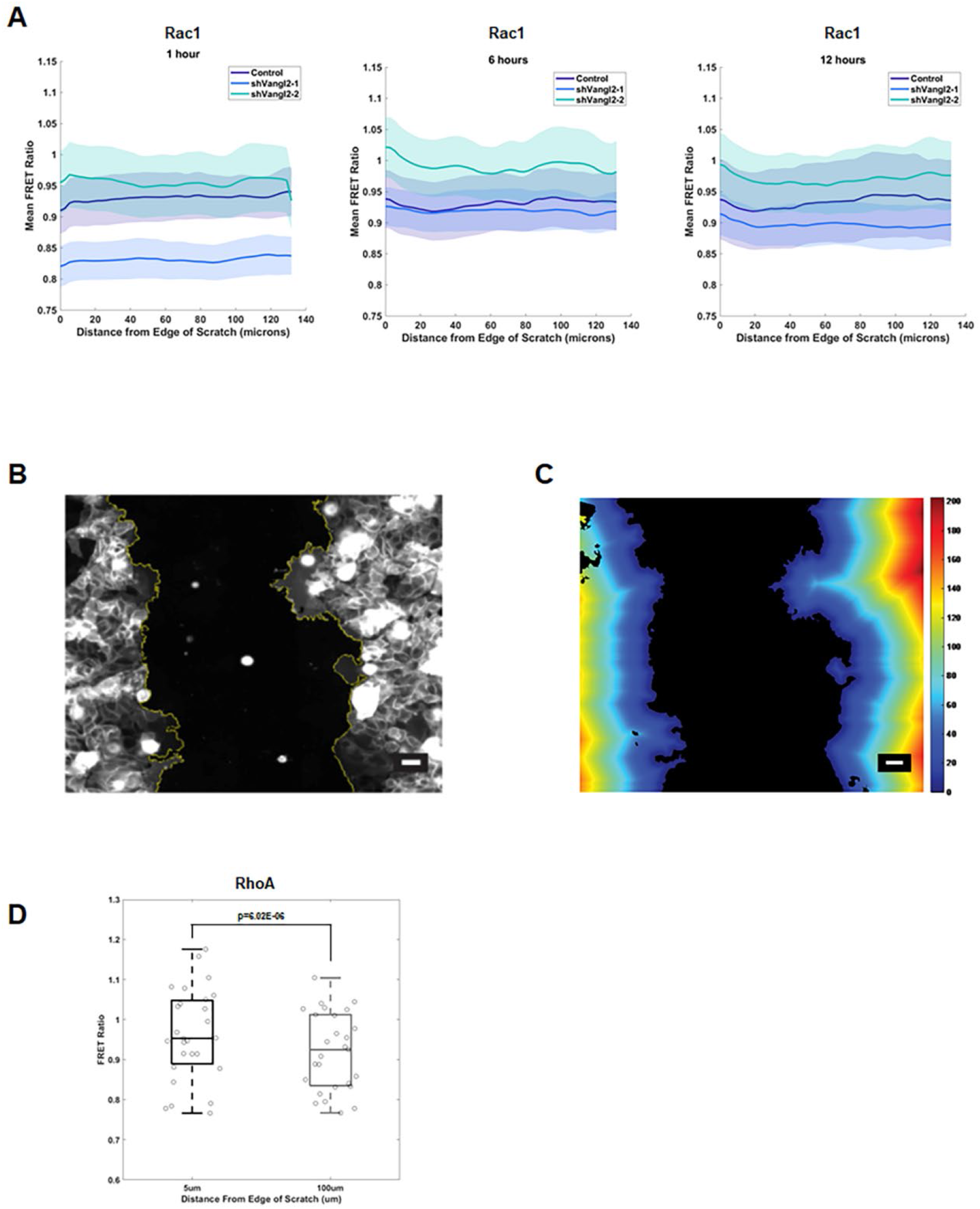
Supporting materials for Figure 6. **A** Rac1 activity as a function of distance in microns from the leading-edge of collectively migrating MCF7 cells stably expressing Vector (*n*=27 wells), shVangl2-1 (*n*=24 wells), or shVangl2-2 (*n*=25 wells) at 1, 6, and 12 hours of migration, error bars indicate ± sem. **B,C** Representative example of custom MATLAB script identifying the leading-edge of a migrating cohort of MCF7 cells **(B)** and binned migrating cells based on their distance from the edge of the scratch in microns, where color bar indicates the distance from the leading-edge of the scratch **(C)**, scale bars=25µm. **D** RhoA activity in MCF7 cells stably expressing the RhoA biosensor and Control at 5µm and 100µm from the edge the of scratch (*n*=27, *p*=6.02E-06), significance was determined by a two-sided paired *t-*test.

